# Adaptive Dynamics of Quantitative Traits in a Steadily Changing Environment

**DOI:** 10.1101/2025.09.28.679017

**Authors:** Sakshi Pahujani, Yuna Zhang, Markus G. Stetter, Joachim Krug

## Abstract

Understanding the adaptation of quantitative traits to changing environments is a central challenge in evolutionary biology. However, the precise roles of the speed of environmental change and trait architecture in adaptive dynamics remain unclear. Here, we investigate their roles through an adaptive walk model and individual-based polygenic simulations within a moving-soptimum framework. We introduce a dimensionless parameter, the effective selection strength, to characterize quantitative traits in both models. We show that adapting populations attain a stationary state below a critical speed of environmental change. Properties of the stationary state are compared between the two approaches, and it is shown that the adaptive walk model provides a good approximation for long-term polygenic adaptation. Additionally, in the adaptive walk model, we derive an explicit formula for the logarithmic divergence of the stationary phenotypic gap upon approaching the critical speed. To elucidate population extinction in biologically realistic terms, we define a fitness-threshold-based critical speed and reveal its power-law dependence on the effective selection strength in both models. Finally, results of our polygenic simulations show how the genetic architecture in the stationary regime is shaped by a dynamic equilibrium between the influx of *de novo* mutations and their loss through fixation and extinction constrained by the rate of phenotypic change enforced by the optimum speed. Taken together, our study advances the understanding of polygenic adaptation and provides insights to simplify analytical work and computational simulations for future studies.

## Introduction

An essential goal of evolutionary biology is to elucidate the genetic architecture underlying adaptation to novel environments (Orr 2005). Quantitative traits, which span a wide range of characteristics, including morphological, physiological, life-history, behavioral traits and the risks of chronic diseases (Lynch and Walsh 1998; Sella and Barton 2019), are of great importance in plant and animal breeding as well as human genetics [e.g., plant height (Moles et al. 2009), milk production (Jiang et al. 2014) and type 2 diabetes (Visscher et al. 2017)]. The continuous variation of these traits makes them excellent candidates for studying evolutionary responses to selective environments, as this continuum allows for the detection of subtle shifts in populations over time. Numerous genome-wide association studies have demonstrated that quantitative traits are typically controlled by multiple or a large number of genetic loci (Buniello et al. 2019). Consequently, adaptation in such traits may result from genetic changes at multiple loci, a process known as polygenic adaptation (Pritchard et al. 2010). The highly polygenic nature of quantitative traits, described by the polygenic model or infinitesimal model, results in a Gaussian distribution of phenotypes (Fisher 1919; Barton et al. 2017). This model has been validated by empirical data (Yang et al. 2010; Boyle et al. 2017).

Selection acting on phenotypes is generally categorized into three forms (Lande and Arnold 1983): directional selection (favoring one extreme phenotype), stabilizing selection (favoring an intermediate phenotype) and disruptive selection (favoring both extremes). Stabilizing selection, as a pervasive evolutionary force, tends to eliminate genetic variation. However, the genetic variance of quantitative traits can be maintained at a stable level under a balance among stabilizing selection, mutation and genetic drift (Bürger 2000). This equilibrium genetic variance has been well studied and can be estimated using mathematical approximations, such as the stochastic House-of-Cards approximation (Turelli 1984).

However, the equilibrium state can be disrupted by environmental changes. Since environmental change is ubiquitous in nature, understanding phenotypic adaptation and the underlying genetic architecture in changing environments has become increasingly important, especially in the context of climate change (Hoffmann and Sgrò 2011). This urgency stems from the escalating impacts of atmospheric *CO*_2_ enrichment, global warming and rising sea levels, requiring systematic study of adaptive response to environmental gradients (Radchuk *et al*. 2019). Populations subject to such changing environments experience directional selection, in addition to stabilizing selection. The directional selection is a consequence of the optimum of the fitness function traversing the phenotypic space due to environmental change (Pease *et al*. 1989). Stabilizing selection when coupled with directional selection, yields a class of moving opti-mum models (Lynch et al. 1991; Lynch and Lande 1993; Bürger and Lynch 1995).

Early quantitative genetic studies in the moving optimum framework mostly utilized the infinitesimal model with adaptation proceeding via the available background standing genetic variation. The assumptions of infinitesimally small effects and a large number of contributing loci allow these models to consider the genetic variation to be a constant (Bürger and Lynch 1995). Such models have offered robust predictions for the response to selection at short time scales (Hill 2014). However, in the more realistic case in which allelic effect sizes vary, significantly different dynamics have been reported for the small-effect and large-effect alleles (relative to a scaled mutation rate) in the short time following a rapid change of the phenotypic optimum (De Vladar and Barton 2014; Jain and Stephan 2017). Similar studies in the context of gradual optimum shifts have underscored the importance of mutations (and genetic architecture) in determining adaptive dynamics (Jain and Devi 2018). To incorporate these increasingly complex and realistic scenarios, forward-in-time simulation has become a powerful and efficient approach to study explicit evolutionary processes and the effects of genetic factors (Carvajal-Rodríguez 2010; Yuan et al. 2012; Messer 2013).

Investigations of *de novo* mutations as the drivers of adaptation have been conducted in parallel, in adaptive walk models (Gillespie 1984; Orr 1998). These models were originally developed in the context of Fisher’s geometric model (Fisher 1930) and rely mainly on the assumption of selection being strong relative to mutation. This renders the adapting population monomorphic at any given time, since a new beneficial mutation fixes instantly upon origin. Collins *et al*. (2007) and Kopp and Hermisson (2009) extended the adaptive walk approximation to study adaptation to a gradually moving optimum and elaborated on the factors that shape the distribution of adaptive substitutions.

Given a moving optimum framework, a key question is the maximum rate of environmental change that an adapting population can tolerate, a subject that has been explored through both quantitative genetics and adaptive walk models. In the quantitative-genetic study by Bürger and Lynch (1995), a critical speed for a population adapting from standing genetic variation was derived based on the phenotypic gap from the optimum becoming too large. Below this critical speed, the population adapts at the same rate as the optimum moves and beyond it, extinction follows rapidly. In contrast, in the moving-optimum supplemented adaptive-walk model of Kopp and Hermisson (2009), a clear critical speed does not emerge because the model, which focused on the effect sizes of fixed mutations, invoked Haldane’s approximation for the fixation probability (Haldane 1927).

Here, we investigate adaptive walks in a moving optimum framework following an analytical construction similar to the work of Nassar and Pardoux (2019). We revoke Haldane’s approximation and derive an expression for the critical speed similar to that reported by Nassar and Pardoux (2019). In a simplified setting where the distribution of mutational effect sizes is replaced by a single effective mutation size, we provide a detailed description of the transition to the non-adaptive state and precisely quantify how the phenotypic gap diverges on approaching the transition. Since the phenotypic gap increases without bound in the non-adaptive regime, the transition occurs at zero population fitness. A more realistic extinction criterion can be formulated by imposing a positive fitness threshold for population viability (Lynch *et al*. 1991; Gomulkiewicz and Houle 2009), and we derive the corresponding fitness-threshold-based critical speed within the simplified model.

We compare and contrast the adaptive walk analysis to individual-based polygenic simulations in which the population is genetically diverse and the selected phenotype is governed by numerous loci simultaneously (Stetter *et al*. 2018). The polygenic model assumes a diploid, randomly mating population of fixed size with genomic parameters based on the model plant *Arabidopsis thaliana*. The simulations confirm that in a gradually changing environment, populations eventually reach a stationary state, while keeping the genetic variance stable. We examine how this stationary state is influenced by the strength of stabilizing selection, the effect size distribution of new mutations and the rate of environmental change, finding partial agreement with the adaptive walk model but also additional features not covered in the simpler setting.

### Model

We consider a one-dimensional quantitative phenotypic trait with the trait value *z*. A Gaussian fitness function *w*(*z, t*) is defined on the phenotypic space,

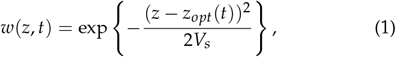

where *z*_*opt*_ (*t*) is the optimum value of the phenotype at time *t* (Figure 1a). The parameter (2*V*_*s*_)^−1^ is inversely proportional to the variance of this function and therefore determines the selection strength. The trait is subject to an environment changing steadily at a speed *v*. As a result, the fitness function and hence the optimum value of the phenotype changes linearly in time according to

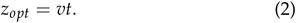

**Figure 1.**
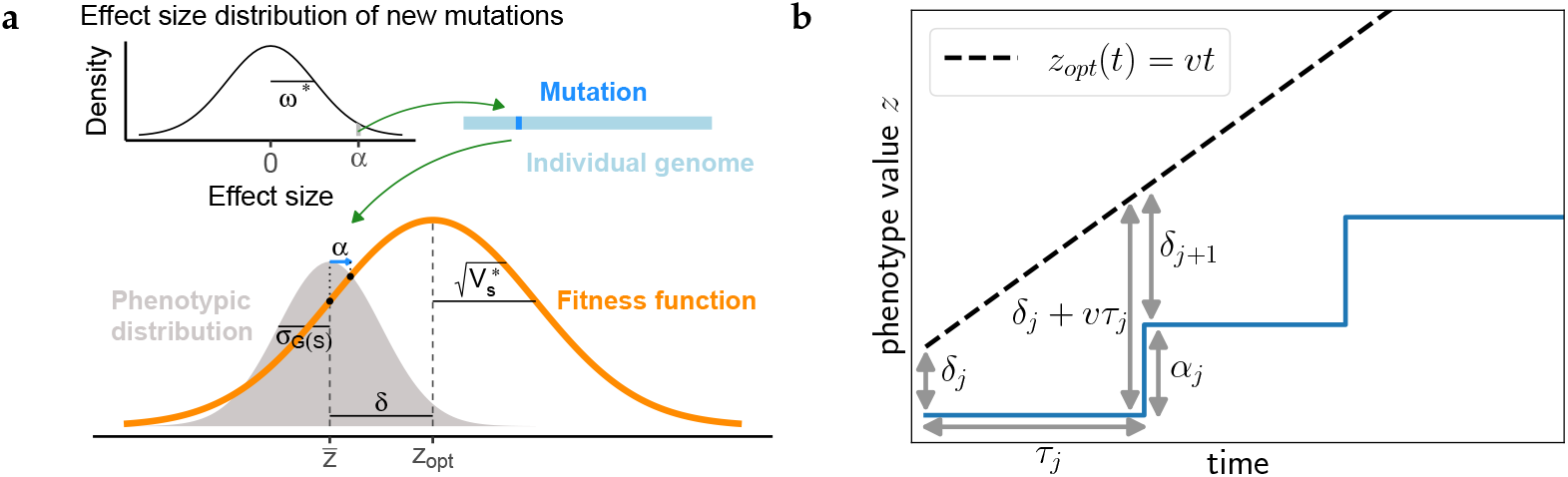
Overview of modeling frameworks. **(a)** The quantitative genetic model of polygenic adaptation. In the bottom panel, the shaded area denotes the phenotypic distribution with a mean 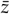 and standard deviation *σ*_*G*(*S*)_. The orange curve shows the fitness function with a peak at the trait optimum *z*_*opt*_ and a width of 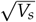, the inverse of which represents the strength of stabilizing selection. The gap *δ* between 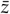 and *z*_*opt*_ indicates how far the current population mean lags behind the optimum. A mutation with a random effect size *α* sampled from a normal distribution 𝒩 (0, *ω*^2^) shown in the top-left plot is introduced into the genome of an individual, resulting in a phenotypic change (*z* → *z* + *α*) and a corresponding fitness change (*w*(*z*) → *w*(*z* + *α*), denoted by two black dots on the orange curve). Variables such as *ω, V*_*s*_ and *v* (with an asterisk) are set to varying values to explore the determinants of polygenic adaptation. **(b)** Adaptive walk model. All individuals of the monomorphic population have the same phenotype value *z*_*j*_ which evolves in a step-like stochastic process.

**Table 1.**
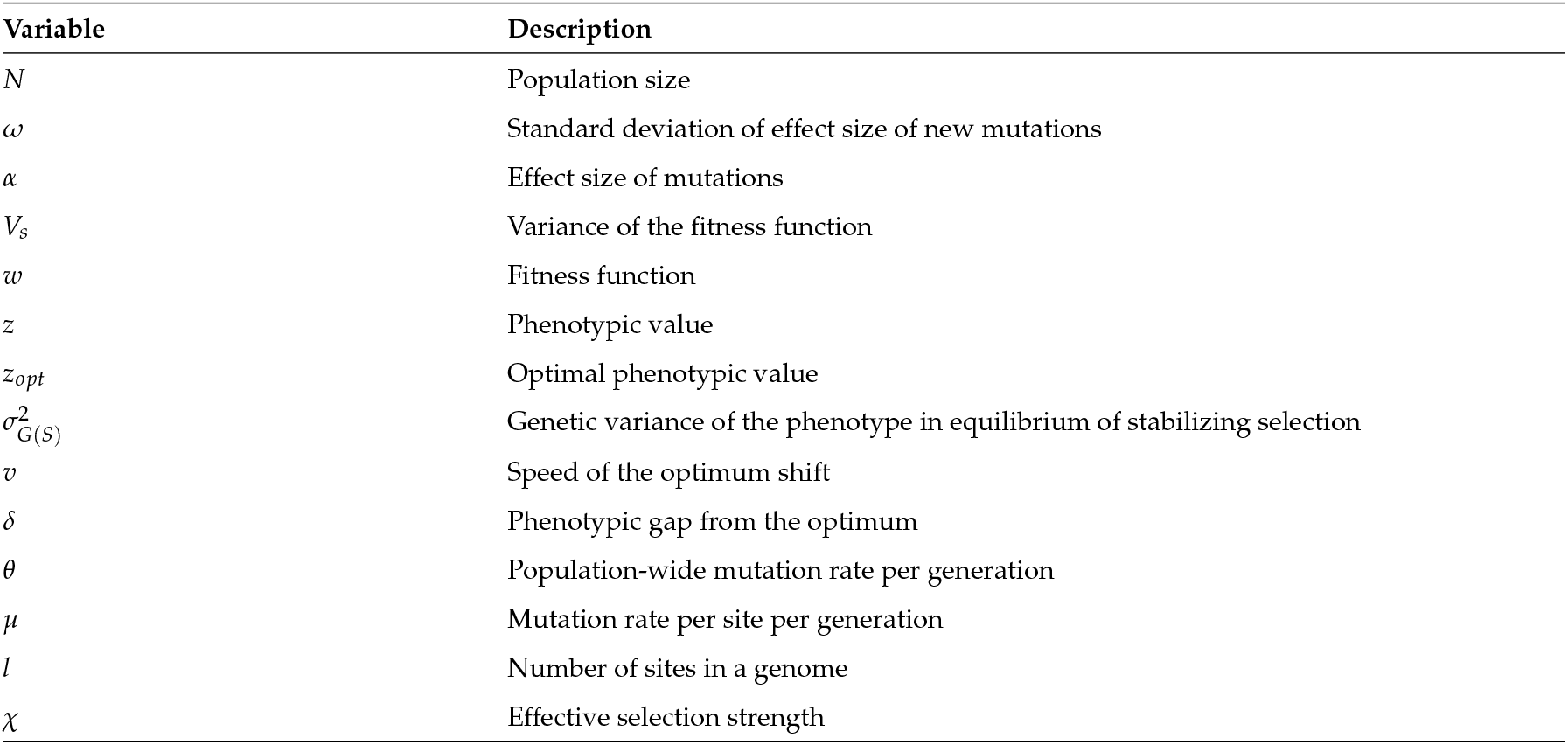
Parameters and variables.

| Variable | Description |
| --- | --- |
| $N$ | Population size |
| $\omega$ | Standard deviation of effect size of new mutations |
| $\alpha$ | Effect size of mutations |
| $V_s$ | Variance of the fitness function |
| $w$ | Fitness function |
| $z$ | Phenotypic value |
| $z_{opt}$ | Optimal phenotypic value |
| $\sigma_{G(s)}^2$ | Genetic variance of the phenotype in equilibrium of stabilizing selection |
| $v$ | Speed of the optimum shift |
| $\delta$ | Phenotypic gap from the optimum |
| $\theta$ | Population-wide mutation rate per generation |
| $\mu$ | Mutation rate per site per generation |
| $l$ | Number of sites in a genome |
| $\chi$ | Effective selection strength |

The population experiencing the changing environment must adapt by introducing new beneficial mutations. Mutations occur at a rate *θ* per generation in the population. The effect-size, *α*, of the incoming mutations is distributed according to a Gaussian with variance *ω*^2^, hereafter referred to as the ‘supply distribution’ or simply the ‘supply’ and defined by

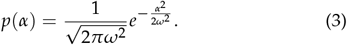

By virtue of the described setting, there are two implicit phenotypic scales in the problem. One is the standard deviation of the fitness function, 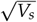 and the other is the standard deviation of the supply, *ω*. In this work, we use the latter to scale the phenotype and all related observables. The ratio of these two scales defines a dimensionless constant

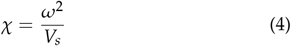

which we refer to as ‘effective selection strength’ or simply ‘selection strength’ in the following. This ratio compares the strides towards the optimum to the imposed strength of stabilising selection, therefore acting as a key determinant of the adaptive process. Another dimensionless parameter is obtained by comparing the speed scales in the problem. The first of these is the speed *v* of the moving optimum. The second is the pace of the adapting population that is set by the phenotypic scale of the effect sizes of mutations and their rate of occurrence in the population. The ratio between the two

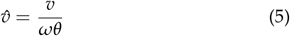

dictates the ability of the population to catch up to the optimum as we will shortly see.

## Methods

### Adaptive walks

To study the model analytically, we restrict ourselves to the strong-selection-weak-mutation (SSWM) regime (Gillespie 1984). This is a regime characterised by the conditions *θ* ≪ 1 (weak mutation) and *s*| *N*| ≫ 1 (strong selection), where *N* is the size of the population and *s* is the selection coefficient that describes the fitness advantage of a mutation. Because of the strong selection, a mutant with a positive selection coefficient can fixate rapidly and one with a negative selection coefficient will be eliminated. Simultaneously, the weak mutation assumption implies that the waiting time between the appearance of two consecutive mutations is much larger than the timescale of fixation. The segregation of two different mutant alleles at the same time in a population under this regime is then highly unlikely, which makes the population monomorphic. The result is a simplified adaptive process with a monomorphic population performing an ‘adaptive-walk’ (Orr 1998, 2005) characterised by periods of waiting times interspersed with instantaneous fixation events. Since phenotypic variability is absent, the state of the population in the SSWM regime is fully described by a single trait value *z*(*t*). The selection coefficient corresponding to a mutation of effect-size *α* is then given by

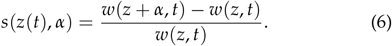

Using the phenotypic gap from the optimum at time *t* defined as

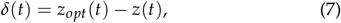

one can rewrite Equation (6) to obtain the selection coefficient as a function of the phenotypic gap,

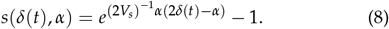

Introducing the standard Gaussian random variable 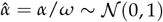 and using the definition of Equation (4), this can be expressed as

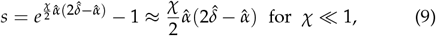

where 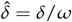 is the phenotypic gap in units of *ω*. In this sense, the dimensionless parameter *χ* sets the scale of the selection coefficients. The parabolic approximation for the selection coefficient stated in Equation (9) is used in the mathematical analysis in the sections that follow, while the exact form has been employed in all simulations.

Within this framework, the phenotypic evolution proceeds in a series of step-like events, which we describe mathematically as a discrete stochastic process. Specifically, we track the gap from the optimum at every occurrence of a mutation^1^. Figure 1b depicts an adaptive walk along with all the associated variables.

The change in the gap at the *j*^*th*^ mutation, *δ*_*j*_, to that at the (*j* + 1)^*th*^ mutation, *δ*_*j*+1_, is written as

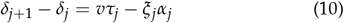

where *τ*_*j*_, *α*_*j*_ and *ξ*_*j*_ are random variables distributed according to

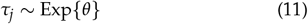

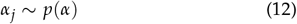

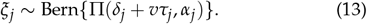

Here, Π(*δ*_*j*_ + *vτ*_*j*_, *α*_*j*_) is the fixation probability of a mutation of effect-size *α*_*j*_ at an instance when the gap is *δ*_*j*_ + *vτ*_*j*_. This probability is given by the large *N* limit of the Kimura formula (Kimura 1962)

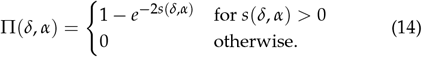

Equation (10) is used to simulate all the adaptive walks described in this work. The first term on the RHS of Equation (10) describes the linear increase in the gap during the waiting period of the (*j* + 1)^*th*^ mutation. Mutations in the model are described as a Poisson process, which implies that the waiting times are exponentially distributed with the mutation rate *θ*. The second term describes the reduction in phenotypic gap by the effect size *α*_*j*_, sampled from the supply, in case of a fixation event occurring with probability Π(*δ*_*j*_ + *vτ*_*j*_, *α*_*j*_). A Bernoulli distributed indicator random variable *ξ*_*j*_, which is 1 with probability Π(*δ*_*j*_ + *vτ*_*j*_, *α*_*j*_) and 0 with probability 1 − Π(*δ*_*j*_ + *vτ*_*j*_, *α*_*j*_), is employed to implement this.

### Polygenic simulations

We employed individual-based simulations using SLiM 4 (Haller and Messer 2023) to generalize the adaptive walk model by incorporating an explicit genotype-phenotype map and multiple segregating mutations. Specifically, we consider a diploid, randomly mating population with a constant size of *N* = 10, 000, where generations are discrete and non-overlapping. The quantitative trait was controlled by a large number of loci within a genome of length *l* = 1 Mb. We used a mutation rate of *µ* = 7 × 10^−10^ per site per generation, which corresponds to 10% of the spontaneous mutation rate observed in the model plant *A. thaliana* (Ossowski *et al*. 2010). Hence, we assumed that 10% of mutations are functional and only simulated functional mutations. This yields a total mutation rate of *θ* = 2*Nlµ* = 14 per generation. The recombination rate was set to 5 × 10^−8^ per base pair per generation, within the range of different plant species (Brazier and Glémin 2022; Salomé *et al*. 2012). An individual’s phenotype was determined solely by the additive genetic effects of all mutations, excluding dominance, epistatic and non-genetic effects. The effect size of mutation, *α*, is randomly drawn from a Gaussian supply distribution with the probability given in Equation (3).

We simulated equilibrium populations under stabilizing selection where the populations evolve in a static environment with a trait optimum of zero for 10 *N* generations (Haller *et al*. 2019). These populations serve as references for adaptive changes resulting from subsequent environmental shifts. To study the moving optimum, we simulated an additional 10 *N* generations, in which the populations evolve to constant optimum shifts as described by Equation (2). The simulation terminated when the fitness of all individuals reached 0 (*<* 10^−300^), which we refer to as “extinction” unless otherwise stated.

To explore the effects of trait architecture on polygenic adaptation, we simulated various quantitative traits by adjusting the strength of stabilizing selection (*V*_*s*_) and the standard deviation of the supply distribution (*ω*). We set *V*_*s*_ to values of 1, 10 and 100, corresponding to a gradient of stabilizing selection from strong to weak, and considered values of *ω* of 0.05, 0.1 and 0.5, representing probabilities for the emergence of large-effect mutations ranging from low to high. Among the 9 combinations of these parameters, two cases (*V*_*s*_ = 1, *ω* = 0.05 and *V*_*s*_ = 100, *ω* = 0.5) yield identical values of the effective selection strength *χ*. For each parameter combination, 100 simulations were run.

Population means and variances in trait values and fitness were recorded for every generation during the 10 *N* generation period of the optimum shift. Additionally, complete population information, including individual phenotypes, fitness values, mutation effect sizes and allelic frequencies, was collected every 500 generations, except during the first and last 1000 generations of the moving-optimum period, where recordings were made every 10 generations. These recordings were used to analyze genetic changes and phenotypic changes during optimum shifts.

## Results

### Critical speed

We first address the adaptability prospects given the rate of change of the environment. Two distinct regimes in the parameter space can be distinguished. In one regime, the population achieves a dynamic stationarity with respect to the moving optimum. In the other, the non-stationary regime or transient regime identified in Nassar and Pardoux (2017), the population cannot keep up with the environmental change and the population mean phenotype diverges from the trait optimum. At the same time, the population fitness is driven to zero. To probe the transition between these regimes as a function of the speed of the optimum, we study the behavior of the phenotypic gap at large speed.

In the non-stationary regime, the gap is expected to be large and we therefore take the limit *δ* → ∞ in the Equation (14) for the fixation probability. In this limit, the selection coefficient (Equation (6)) is positive and large for all mutations that increase the phenotypic value, *α >* 0, and correspondingly

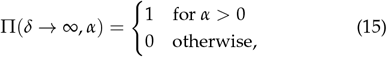

i.e., every mutation sampled from the supply is fixed as long as the effect size is positive. The distribution of the fixed effect sizes takes the form of a half-Gaussian on the set of positive effects, and the mutation rate is simultaneously reduced to half of the original value. The rate at which the population adapts is therefore 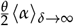, where

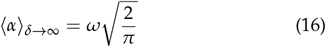

is the mean of the half-Gaussian. This is the maximally possible rate of adaptation, which defines the critical speed *v*_*crit*_. In dimensionless units, it reads

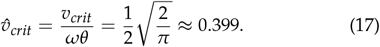

Adaptive walk simulations confirm the prediction in Equation (17). The mean phenotypic gap converges for speeds below the critical value, and increases without bound above the critical speed (Figure 2a). This behaviour is consistent across all values of the effective selection strength *χ* that we simulated.

**Figure 2.**
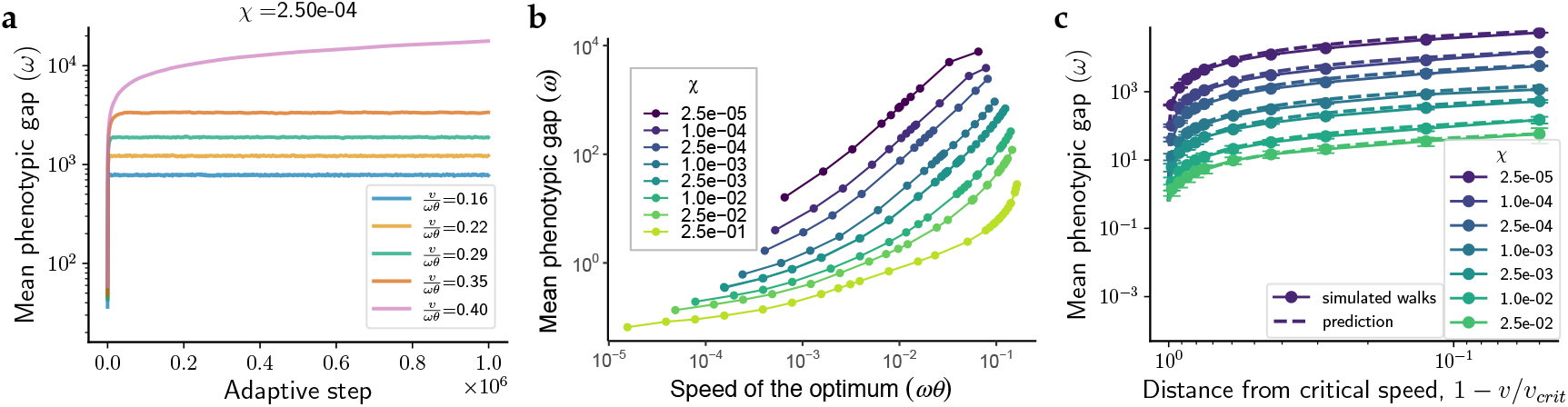
The phenotypic gap. **(a)** The critical speed. The evolution of the mean phenotypic gap over the course of simulated adaptive walks is depicted at speeds of the optimum up to the critical speed. Each curve is an average of the phenotypic gaps recorded from 100 independent walks just before an adaptive step. With *ω* and *θ* set to 1, the critical speed here is 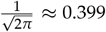. For speeds below this critical value (stationary regime), the mean phenotypic gap can be seen to converge whereas at the critical speed, it increases indefinitely (non-stationary regime). **(b)** The mean phenotypic gap over the last 1000 generations from 100 independent polygenic simulations. The speed of the optimum shift is in units of *ωθ* and the phenotypic gap is in units of *ω*. **(c)** Stationary phenotypic gap as a function of the distance to the critical speed in the adaptive walk model. The scattered points are obtained from simulations using a Gaussian supply and the exact form Equation (6) of the selection coefficient. The dashed lines are theoretical predictions using the effective mutation size approximation. Note that, in this representation, the speed of the optimum increases from left to right and the critical speed is located at positive infinity. Figure S1 shows the same data as a function of 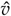.

Consistent with the adaptive-walk model, polygenic simulations reveal that the phenotypic gap reaches a stationary state when the speed of the optimum shift remains below a certain threshold (Figure S5). However, when the shift speed exceeds this threshold, the phenotypic gap continuously widens, eventually leading to population extinction (Figure S6). To understand the non-equilibrium stationary state that populations establish in changing environments, we analyzed the stationary values (operationally defined as averaged values over the last 1000 generations) of phenotypic gaps and other observables. Our results demonstrate that both the effective selection strength and the speed of the optimum shift significantly influence the stationary phenotypic gap. Specifically, weaker effective selection strengths produce larger stationary gaps when the trait optimum shifts at equal speeds. Similarly, higher shift speeds result in larger stationary gaps across all effective selection strengths (Figure 2b). These trends are in qualitative agreement with the adaptive walk model (Figure 2c and Figure S1) and will be explained in detail in Section **Effective mutation size approximation**.

### Fixed mutations

Phenotypic adaptation is driven by changes in allelic frequencies. Positive-effect alleles are favored when the mean phenotypic value lags behind the trait optimum, and thus their frequencies increase. This can eventually result in the fixation of mutations if the selection pressure is strong enough. We found that the number of fixation events per generation (*n_f_*) and the mean effect size of mutations that fixed 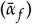 become stationary over the course of polygenic simulations (Figures S7 and S8). This adaptive dynamic satisfies the constraint

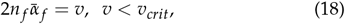

where the left-hand side represents the rate of phenotypic changes due to fixed mutations and the factor 2 is specific to the diploid setting of the simulations (Figure S19). Therefore, populations maintain persistent adaptation by tracking the moving phenotypic optimum through successive fixation processes. This indicates that the adaptive walk model is a good approximation for studying the adaptive dynamics of polygenic traits in the long term.

In addition, the stationary values of *n_f_* and 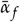 exhibit pre-dominant dependence on the speed of optimum shift but remarkably weak modulation by the effective selection strength *χ*. More specifically, the number of fixation events *n_f_* is linearly related to the speed of the optimum shift, yet independent of *χ* in both the adaptive walks and the polygenic case (Figures S2 and 3a). Similarly, the stationary 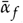 shows little (for lower speeds) or no variation (for higher speeds) with *χ* in both models (Figures 3b and 3c). Unexpectedly, the dependence on the speed of the optimum is found to be non-monotonic. This property can be attributed to the interplay of the supply and the fixation probability and is explained in the Appendix.

**Figure 3.**
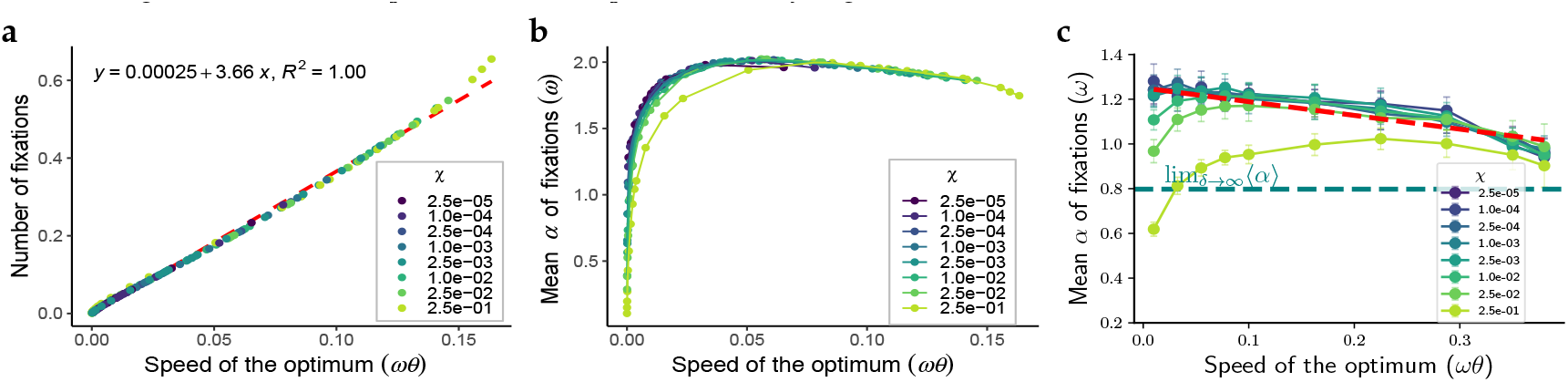
Properties of fixed mutations. **(a)** Number of fixation events per generation. **(b)** Mean effect size of fixed mutations in units of *ω*. The results in panels **(a-b)** are averages over the last 1000 generations of 100 independent polygenic simulations. The number of fixation events in the adaptive walk model is shown in Figure S2. **(c)** Mean effect size of fixed mutations in the adaptive walk model. For small values of the effective selection strength *χ* the data converge to a single curve which decreases approximately linearly with increasing optimum speed. The dashed line shows a linear fit to these curves, which defines the effective mutation size 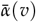.

### Effective mutation size approximation

As was shown above, the mean effect size of fixed mutations is only weakly dependent on *χ* and for the most part, the dependence on the speed of the optimum can be considered linear in the adaptive walk model (Figure 3c). We exploit this property in favor of simplifying the problem by introducing the effective mutation size *α*(*v*) as a linear function of the speed of the optimum shift. Replacing the random variable *α*_*j*_ in Equation (10) by a constant *α* eliminates the nonlinearity and enables us to explicitly average the right-hand side with respect to the remaining random variable *τ*_*j*_. Denoting by Δ*δ* = ⟨*δ*_*j*+1_ − *δ*_*j*_⟩ the average change in the phenotypic gap, this yields

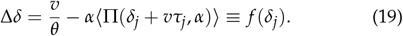

The right-hand side of the equation is a function *f* (*δ*) of the initial gap *δ*_*j*_, which is derived in the Appendix and depicted in Figure 4a.

**Figure 4.**
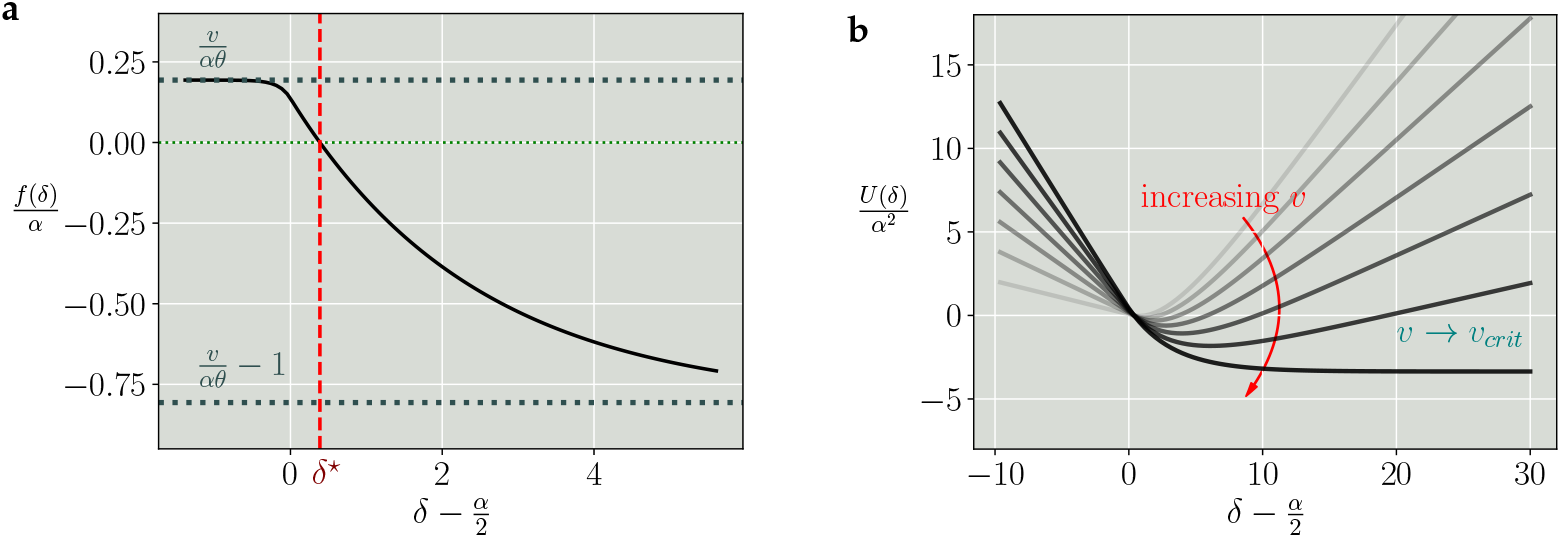
Force and potential governing the change of the phenotypic gap. **(a)** The force *f* (*δ*) defined in Equation (19) is plotted for parameters in the stationary regime. For 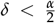, the force is a constant. In this case the gaps are smaller than the minimum value required to yield a positive selection coefficient for the fixed *α*. The force stems from the average waiting time and drives an increase in the gap. Once the gap becomes large enough such that the selection coefficient is positive, the force decreases and eventually asymptotes to the limiting value of 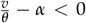. The force crosses zero at *δ* = *δ*^⋆^. **(b)** The potential, *U*(*δ*) is plotted as a function of 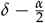 for varying speeds of the optimum. For speeds below the critical speed, the potential is confining, with a minimum at *δ*^⋆^. As the speed hits the critical value, the right arm of the potential flattens and it loses its confining property.

Equation (19) suggests to interpret the function *f* (*δ*) as a force driving the change in the phenotypic gap: the gap increases when *f >* 0, decreases when *f <* 0 and remains stationary at the value *δ*^*^ defined by *f* (*δ*^*^) = 0. Moreover, to elucidate the transition to the non-stationary regime, it is instructive to introduce a potential *U*(*δ*) related to the force through

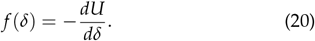

In this way, the time evolution of the gap can be visualized as the motion of a particle driven by random fluctuations and confined by the potential *U*(*δ*). This viewpoint has been adopted in previous work on the adaptive walk problem working in small-jumps/weak selection limit. In this limit, the confining potential is harmonic and the stochastic process governing the dynamics of the gap is known as an Ornstein-Uhlenbeck process (Kopp et al. 2018; Nassar and Pardoux 2019; de Souza Silva et al. 2023).

The potential *U*(*δ*) is a U-shaped function of the gap and has a right arm that features a decreasing slope with increasing speed of the optimum. As the speed reaches the critical value, the right arm of the potential becomes completely flat (Figure 4b). This property of the potential provides a precise description of the transition from the stationary to the non-stationary regime. In the stationary regime, the potential is confining, which keeps the phenotypic gap finite. At the critical speed, the potential is no longer confining and the gap diverges. In the stationary regime, the potential has a unique minimum for a given value of the speed. The minimum can be found by equating the force *f* (*δ*) to zero. It represents the stationary value of the mean gap from the optimum and is given by

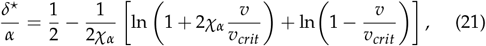

where 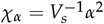 is the effective selection strength and *v*_*crit*_ = *αθ* is the critical speed in the effective mutation size formulation of the adaptive walk model.

For speeds close to the critical speed, Equation (21) is dominated by the second term in the square brackets. In the limit *v* → *v*_*crit*_, the stationary gap diverges logarithmically with a prefactor that is inversely proportional to the effective selection strength *χ*_*α*_. The logarithmic increase is very gradual, which implies that the eventual divergence of the gap at *v*_*crit*_ cannot be anticipated even very close to the critical speed (Figure S1). Finally, we compared the prediction for *δ*^⋆^ from Equation (21) obtained from the effective mutation size approximation to the stationary value of the mean gap obtained from adaptive walk simulations that draw the mutation effect sizes from a Gaussian supply distribution and found an excellent agreement of these two (Figure 2c).

### Fitness threshold for extinction

So far, the critical speed *v*_*crit*_ separating the adaptive from the non-adaptive regime was defined as the value at which the stationarity of the gap in the adaptive walk model is lost. While this criterion is precise and mathematically appealing, it is biologically unreasonable because it assumes that the population remains viable at arbitrarily low fitness. A more realistic criterion can be obtained by introducing a fitness threshold *w*_*th*_ *>* 0 for viable populations (Lynch et al. 1991; Gomulkiewicz and Houle 2009; Matuszewski et al. 2014). In polygenic simulations, population mean fitness decreases as the speed of optimum shift increases. Notably, for stronger selection, the fitness profiles are seen to shift uniformly to the right, which implies that the threshold speed *v*_*th*_ at which the fitness threshold is reached increases with increasing *χ* (Figure 5a).

**Figure 5.**
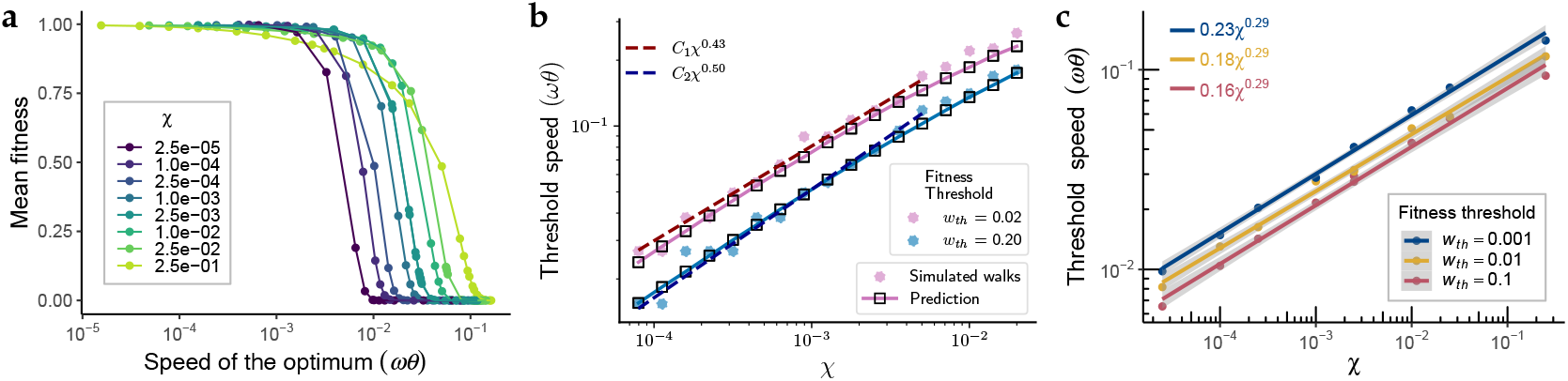
The threshold speed. **(a)** Mean fitness over the last 1000 generations of 100 independent polygenic simulations. **(b)** The threshold speeds acquired from adaptive walk simulations are plotted against the prediction (Equation (C.5)) obtained from the effective mutation size approximation. In the regime *χ* ≪ 1 shown here, Equation (C.5) reduces to the power law (Equation (23)). **(c)** Threshold speeds obtained at three fitness cutoffs (indicated by colors) from polygenic simulations. The equations of the fitted lines are shown in the upper left corner. In panels **(b)** and **(c)**, both axes are shown on logarithmic scales.

Within the effective mutation size approximation, the threshold speed *v*_*th*_ can be obtained by determining the value of the stationary phenotypic gap 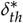 at the threshold from the relation

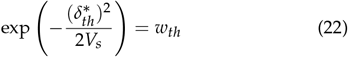

and inverting Equation (21). We compared the analytic expression for the threshold speed (Equation (C.5)) to adaptive walk simulations using the full Gaussian supply distribution and found good agreement between them (Figure 5b). For small values of the effective selection strength, Equation (C.5) reduces to a square-root dependence

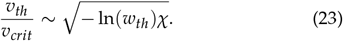

In the polygenic simulations, we found a similar power-law relationship between the threshold speed and the effective selection strength, expressed as 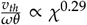 with a prefactor that depends weakly on *w*_*th*_ (Figure 5c). We also scaled the speed by the square root of the equilibrium genetic variance, and found a similar power-law relationship with different exponents around 0.7 (Figure S10 and Appendix). Thus, for both adaptive walks and polygenic simulations, the threshold speed obeys power-law scaling with the selection strength, but with different leading exponents.

### Genetic variance

Since the adaptive walk model assumes a monomorphic population, we extended our work by polygenic simulations to obtain insights pertaining to the genetic variance and changes in genetic architecture. Prior to the environmental change, populations are in an equilibrium state maintained by the balance between mutation, stabilizing selection and genetic drift. This state is characterized by a stable level of genetic variance (Figure S11). The environmental change disrupts this balance, altering the genetic variance. We found that genetic variance initially increases during slow optimum shifts or decreases during fast optimum shifts. In both cases, the variance subsequently stabilizes at a new level (Figure S12). This shows that a new, non-equilibrium stationary state is established even as the optimum continues to move. In the following, we describe some of the statistical properties of this state, beginning with the amount of genetic variance.

The stationary level of genetic variance is influenced by both the shift speed and the selection strength. At large selection strength, corresponding to low equilibrium genetic variance, the stationary non-equilibrium genetic variance increases with shift speed, but this growth is not unlimited. Beyond a certain speed of the optimum shift, the stationary genetic variance begins to decrease. The inflection points, where the trend shifts from increasing to decreasing, occur at higher shift speeds for populations experiencing stronger selection strengths (Figure 6b). Furthermore, the influence of the effective selection strength on the stationary genetic variance is most pronounced at slower shift speeds. For a given shift speed, traits with higher values of *χ* exhibit lower genetic variance. However, as the shift speed approaches the critical speed, genetic variance converges across all examined *χ*, indicating that the influence of the strength of stabilizing selection 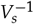 becomes negligible under these conditions (Figure 6b).

**Figure 6.**
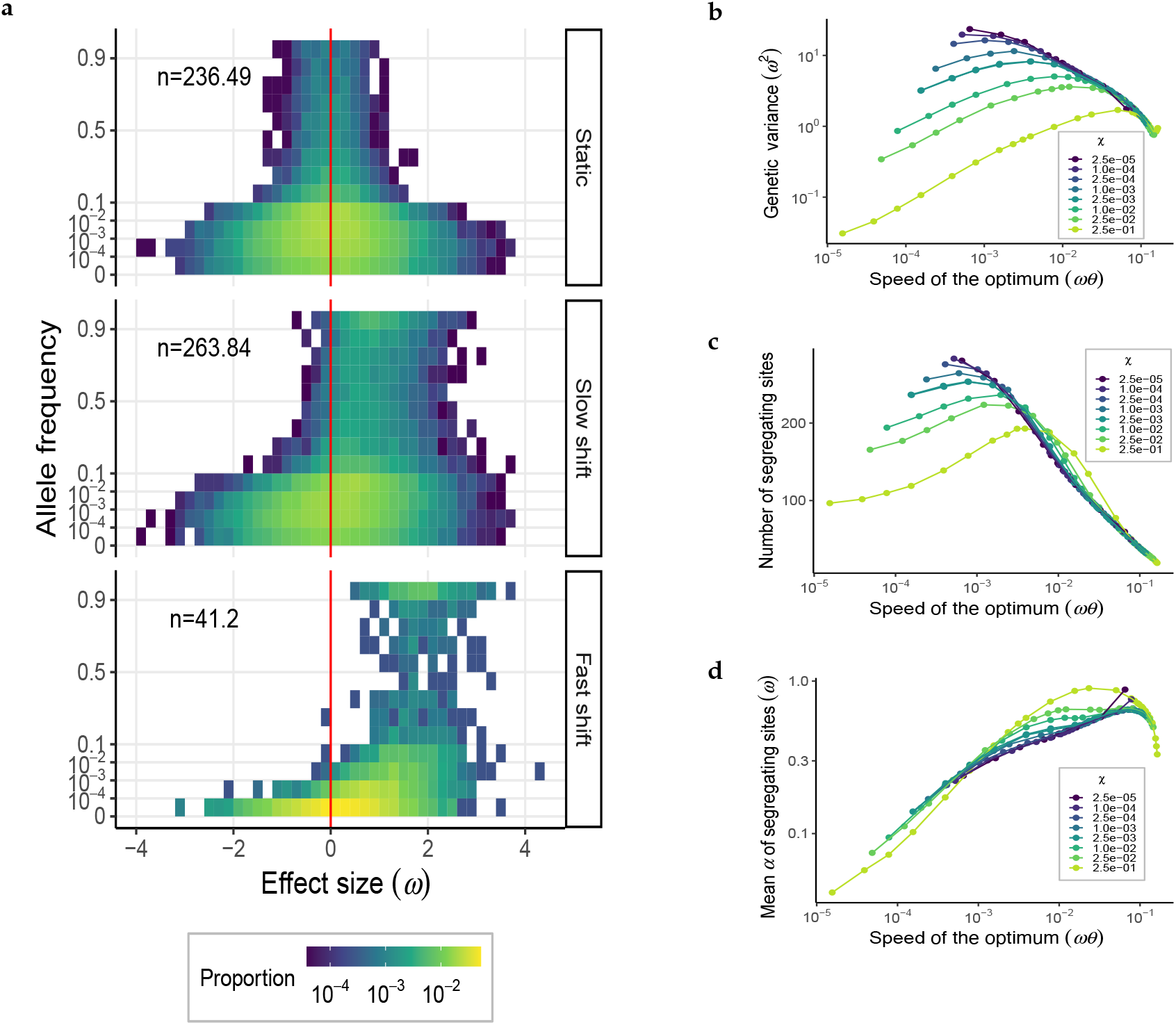
Genetic basis of polygenic traits. **(a)** The genetic architecture matrices for effective selection strength *χ* = 10^−3^ under the conditions of static, slow (6 × 10^−4^ *ωθ*) and fast (9.6 × 10^−2^ *ωθ*) optimum shift. The red line separates positive and negative effect sizes. The proportion of segregating sites within a certain range of effect sizes (in units of *ω*) and allelic frequencies is indicated by color. The number in the upper left corner denotes the total amount of segregating sites in the population, averaged over 100 replicates. **(b)** The mean genetic variance which is in units of *ω*^2^. **(c)** The mean number of segregating sites per generation. **(d)** The mean effect size of segregating sites. In **(b-d)**, the results are averaged over the last 1000 generations of 100 independent polygenic simulations.

### Genetic architecture during adaptation

Apart from its effect on genetic variance, the environmental change shapes the genetic architecture underlying adaptive traits in other ways. Compared to the equilibrium distribution of effect sizes before the optimum shift, there are more positive-effect alleles with higher frequencies and fewer negative-effect alleles in the population. During a slow environmental shift, the total number of segregating sites increases compared to that before the shift, while fast shifts led to the opposite result (Figures 6a and S15 to S17).

To explain the non-monotonic dependence of the number of segregating sites on the shift speed, note that many positive-effect alleles are favored by selection during optimum shifts and thus maintained in the population. On the other hand, environmental shifts lead to the fixation or loss of alleles, which reduces the number of segregating sites. Since the number of fixed mutations is independent of the effective selection strength *χ* (Figure 3a), the difference in the number of segregating sites across different values of *χ*, where populations with greater *χ* retain fewer segregating sites under the same shift speed, suggests that more maladaptive mutations are purged under stronger selection strengths. However, this difference diminishes at higher shift speeds, particularly when approaching the critical speed (Figure 6c). In this regime, the number of segregating sites is a decreasing function of the shift speed that is independent of *χ*.

The mean effect size of segregating sites generally increases with shift speed but declines as the speed approaches the critical speed – a pattern similar to that for fixed mutations (Figure 6d, see the comparison in Figure S18). This trend is attributed to the fact that as the speed of environmental shift increases, more positive-effect mutations increase in frequency, while negative-effect mutations tend to decrease their frequencies or are lost, leading to an increase in mean effect size. However, when the speed approaches the critical speed, many large-positive-effect mutations become fixed, which consequently reduces the average.

## Discussion

### Adaptive regimes and the critical speed

In the face of a steadily changing environment, a population experiences directional selection in addition to the stabilizing selection present at equilibrium. Such directional selection can be represented by a fitness function traversing the phenotypic space (Pease *et al*. 1989). The population must respond adequately to the directional selection in order to avoid extinction. Since natural populations are subject to genetic constraints (Barton and Partridge 2000), an upper bound on the tolerated speed of environmental change limits the adaptive potential.

A critical speed was derived for adaptation from standing variation already in the early 1990s (Bürger and Lynch 1995). However, the concept has only recently been explicitly discussed for long-term adaptation from *de novo* mutations. Within the adaptive walk framework, Nassar and Pardoux (2017) identified a recurrent regime and a transient regime separated by a critical speed. The transient regime was absent in earlier studies of the adaptive walk model, because they employed an approximate expression for the fixation probability that did not saturate at large selection coefficients (Kopp and Hermisson 2009). Here we describe the two regimes as stationary and non-stationary, respectively. In the stationary regime, the phenotypic gap of the adapting population from the optimal phenotype attains a finite mean value, while it increases indefinitely in the non-stationary regime. The derived critical speed depends only on the mutation rate and the standard deviation of the supply distribution. It defines the maximum possible speed at which a population can adapt.

Our study includes polygenic simulations, which allowed us to incorporate a more complex genetic make-up of the population and to explicitly examine the associated changes in the genetic architecture during the optimum chase. The simulated populations go extinct much before the predicted critical speed is reached. This discrepancy indicates that the assumption underlying Equation (17) – that all positive-effect mutations can eventually be fixed – is not valid in the polygenic case. In polymorphic populations, a positive-effect mutation can be selectively disadvantageous even if the average phenotype is below the optimum, as long as it occurs in an individual with a phenotypic value greater than the optimum. Additionally, genetic drift and linkage disequilibrium may cause a selectively advantageous mutation to be lost by chance (Stetter *et al*. 2018).

This shows that while the adaptive walk model is a well-suited approximation for the long-term pattern, it cannot capture the competing forces in a polygenic background.

### The steady state of directional selection

Despite the complexity of the evolutionary forces acting in the stationary regime of polygenic adaptation, the resulting steady state can be described rather simply. In particular, the balance relation (Equation (18)) shows that the rate of phenotypic evolution is constrained by the speed of the optimum shift and fully determined by the properties of fixed mutations. This shows that the pool of segregating mutations serves as a stable supplier of fixed mutations while producing an invariant phenotypic effect. Underlying this process, the genetic architecture remains invariant through dual fluxes: a continuous influx of novel mutations and an outflux via fixation or loss of mutations. This dynamic equilibrium manifests itself as temporal stability in the number of segregating loci, the mean effect size, and the mean allele frequency (Figures S13, S14 and S20), resulting in stationary genetic variance under constant environmental change.

### The effective selection strength

The observables characterizing the stationary regime exhibit a crucial dependence on the dimensionless parameter *χ* defined in Equation (4) that we referred to as the effective selection strength. While the selection coefficients of individual mutations depend on the background through the phenotypic value *z*(*t*), see Equation (6), their overall magnitude is determined by *χ*. Intuitively, *χ* compares the mutational advances toward the optimum to the imposed strength of stabilizing selection.

This parameter has been somewhat omnipresent in previous studies, see e.g. Hayward and Sella (2022) and Milligan *et al*. (2025) and references therein. In Kopp and Hermisson (2009), *χ* is embedded within the composite parameter *γ*, which incorporates both genetic and environmental factors that govern the distribution of adaptive substitutions. A higher-dimensional analogue of the effective selection strength appears in the study of multiple pleiotropic traits, where the eigenvalues of the product of the selection and mutation matrices determine the distribution of fitness effects and the effective number of traits under selection (Martin and Lenormand 2006).

In nature, different traits have different distributions of effect sizes and different selection pressures acting on them. Characterizing traits by *χ* allows the study of traits that effectively experience the same strength of selection under the same umbrella. Therefore, the parameter *χ* can be employed to reduce the dimensionality of parameter space in quantitative genetic studies.

Our results identify distinct regimes and observables that are strongly influenced by *χ*, as well as others where its influence diminishes. We find that *χ* maintains a strong influence on observables associated with the adaptability of a population, namely the phenotypic gap and the threshold speed. The reduction of the phenotypic gap observed at higher selection strengths (Figure 2) is an effect that also manifests itself as an increase in the threshold speed (Figure 5). When *χ* is small, even a modest optimum shift leads to a substantial drop in fitness because the means to adapt are restricted. Increasing *χ*, while holding the rate of environmental change constant, initially raises the tolerated rate of change, but further increases in selection strength have diminishing returns. With the sensitivity to the selection strength lowered, the maximum speed tolerated approaches the critical speed *v*_*crit*_, marking the onset of a regime where adaptation is limited not by the lack of sufficient selection pressure (‘selection-limited’), but by the mutation rate and supply, i.e., the genetic factors (Figure S4). Notably, we found that the influence of *χ* does not extend to the number and mean effect size of mutations that fix in the stationary state (Figure 3). Once the phenotypic gap has been established in the early phase of adaptation, it is maintained by fixed mutations in a way that is independent of the selection strength.

The decline in genetic variance follows naturally from stronger purifying selection acting on high-*χ* traits, which leads to increased purging of segregating alleles. After the onset of the genetically limited regime, directional selection dominates, rendering the influence of stabilizing selection negligible. This transition results in the convergence of genetic variance across traits, with the speed of environmental change emerging as the key determinant (Figure 6).

## Conclusion and outlook

The adaptive-walk model integrated with polygenic simulations provides a convenient framework for exploring the adaptation of quantitative traits, yet several caveats deserve attention. Our results rely on an idealized scenario of steady environmental change, while real-world environmental changes are often irregular and unpredictable (Marrec and Bank 2023). In addition, our framework presumes known and constant values for both the distribution of effect sizes of new mutations (*ω*) and the variance of the fitness function (*V*_*s*_), which are challenging to estimate and may vary over time in nature.

Despite these limitations, employing reasonable assumptions for *ω* and *V*_*s*_ in a theoretical construct allowed us to tap into the details of long-term adaptation of quantitative traits. This includes examining the precise behavior of the phenotypic gap, tracking the changes in genetic architecture with the progression of the adaptive process, and studying other observables that may be experimentally inaccessible. Although a direct comparison of these observables with empirical data can be considered ambitious, investigating them theoretically furthers our understanding of the properties of the adaptive process. For instance, the slow divergence of the phenotypic gap upon approaching the critical speed tells us that adaptation is a rather efficient process – the phenotypic gap can be maintained up until the population hits its inherently limited adaptation speed. Additionally, we can extrapolate our findings of the critical speed to accelerating/decelerating environments (Greenspoon and Spencer 2021). Since the critical speed is a property of the adapting population (and not the changing environment), one can expect a stationary regime even under accelerating/decelerating environments up to the point where the environmental change outpaces the critical value. Finally, our in-depth analysis informs which parameters can be most effectively manipulated to facilitate informed decisions.

## Data availability

Code used for this study can be found at https://github.com/YunaZhang73/simulate_CCE.

## Acknowledgments

We thank the members of TRR 341 *Plant Ecological Genetics* for insightful comments and discussions.

## Funding

Funded by the Deutsche Forschungsgemeinschaft (DFG, German Research Foundation) – Project-ID 456082119 – TRR 341/1

## Conflicts of interest

The authors declare no competing interests.

## Appendices

### A. Non-monotonic dependence of the mean effect size on the speed of the optimum

In the adaptive walk picture, the change in the phenotypic gap depends, among other things, on the size of the mutations that fix. The probability density, Ω(*α*| *δ*) for a mutation of effect size *α* to fix in the population with a gap of *δ* from the optimum is proportional to the probability of it being sampled from the supply and the corresponding fixation probability, i.e.,

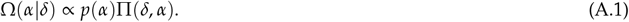

The supply distribution is invariant in the adaptive process and favours small effect mutations. The fixation probability on the other hand, depends on the gap and is positive only when the selection coefficient corresponding to the sampled effect size at the given gap is positive, i.e., 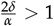. For a finite gap, the maximum of this fixation probability occurs at *α* = *δ*, i.e., for the perfect step that covers the gap exactly. The fixation probability for this step, Π_*max*_ is

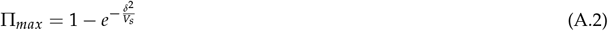

Thus, the size of the perfect step increases with increasing gap and so does the corresponding fixation probability. This indicates that the fixation probability becomes increasingly more biased towards larger step sizes as the gap increases (see Figure S3(a)). Since the supply distribution is independent of the gap, the effect of this bias is seen as an increase in the mean effect size of fixed mutations as the speed of the optimum and hence the gap increases.

The fixation probability, however, is bounded and must therefore stop increasing with increasing gap. This is reflected in the curvature of the fixation probability at its maximum,

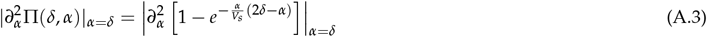

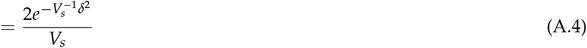

The curvature at the maximum decreases exponentially with *δ*^2^. With increasing gap therefore, the fixation probability tends to a square function of the effect size bounded between *α* = 0 and *α* = 2*δ* (Figure S3(b)). Since most effect sizes in that domain become equally probable for fixation, the bias of the supply distribution to the smaller sizes results in a decrease in the mean fixation size observed in Figure 3c at higher speeds. As *δ* → ∞, the fixation probability Π(*δ, α*) → 1 for step sizes between 0 and 2*δ* and as discussed in the main text, 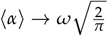 (see Equation (16)).

The fixation probability is also a square function for step sizes between 0 and 2*δ* for a finite *δ* but with 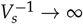. In this case, for a given gap *δ*, the size of a step is sampled effectively from a part of the Gaussian supply that is cut-off at *α* = 0 and 2*δ*. As the gap increases, a larger and larger part of the Gaussian supply is “swept-through” by the fixation probability (Figure S3(c)). Thus, the mean fixation size in this limit increases monotonically. It can in-fact be found as the expectation value of the Gaussian with a finite support and conditioned on the gap.

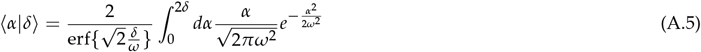

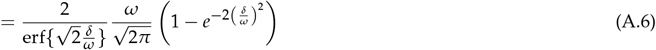

The mean again asymptotes to 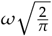 as *δ* → ∞.

### B. Force for fixed effect size

The force, *f* (*δ*) is derived by taking the average of Equation (10) for a fixed mutational effect size *α*_*j*_ ≡ *α*, conditioned on *δ*_*j*_ = *δ*. The indicator random variable *ξ*_*j*_ is given by

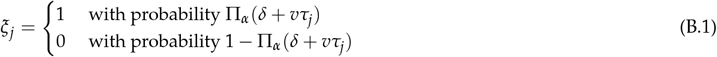

where

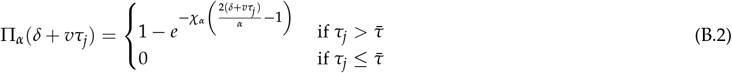

and

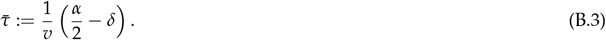

We distinguish two cases:

I. 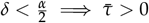
II. 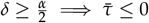

#### Case I: 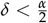

First, we fix *τ*_*j*_ = *τ* and average Equation (10) with respect to the random variable *ξ* which gives^2^:

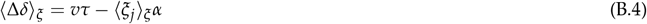

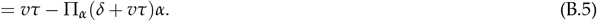

Next, we average with respect to *τ* which yields

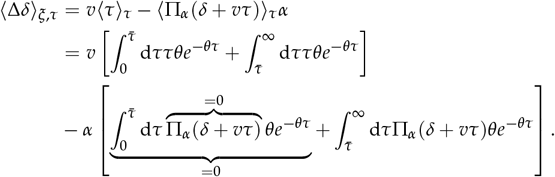

The above evaluates to

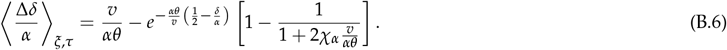

#### Case II: 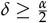

In this case, 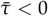. Since, the waiting time is always positive, it is by extension greater than 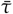. We proceed as before by first fixing *τ*_*j*_ = *τ* and averaging Equation (10) with respect to *ξ*.

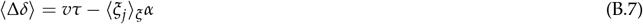

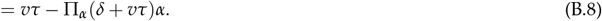

Then, averaging with respect to *τ* gives us

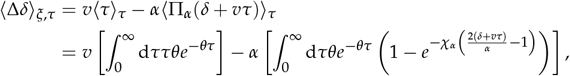

which evaluates to

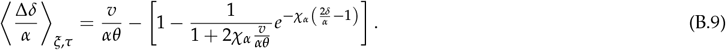

Using *v*_*crit*_ = *αθ* along with Equation (B.6) and Equation (B.9), we obtain

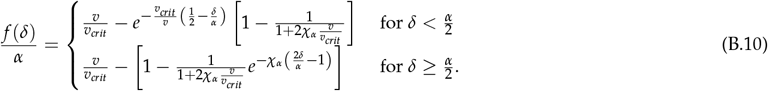

The fixed point, *δ*^⋆^ is found using

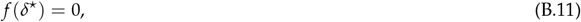

For the first case, 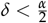, the derived *δ*^⋆^ is larger than 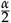 and is therefore invalid. The second case provides a well behaved expression:

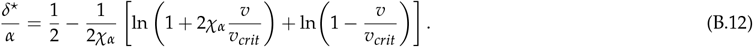

The force can be integrated and yields the following potential, *U*(*δ*):

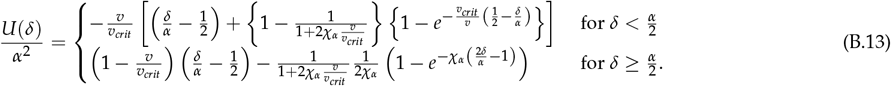

### C. Derivation of the threshold speed

We introduce a fitness threshold *w* = *w*_*th*_ which is attained at a phenotypic gap *δ* = *δ*_*th*_. Solving Equation (22) gives

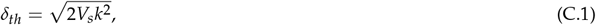

where we have defined *k*^2^ =− ln *w*_*th*_ ≥ 0.

In the following our goal is to find the threshold speed, *v*_*th*_ at which this threshold gap is reached. We work again within the fixed-*α* approximation where the force is given by Equation (B.10). The threshold speed *v*_*th*_ is then the speed that solves

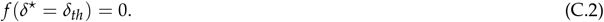

Evaluating the above for 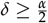 leads to

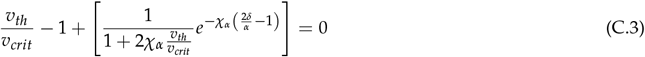

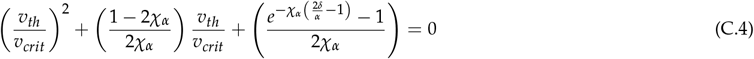

This is a quadratic equation with two solutions. However, the only viable solution is one which yields a positive threshold speed. Using Equation (C.1), we get

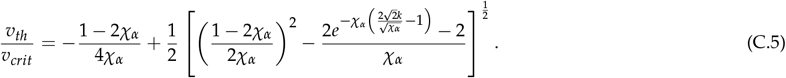

Note that due to Equation (C.1), the imposition 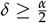 also implies a maximum value of the fitness threshold for which Equation (C.5) is valid. This upper cap is given by 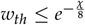 or 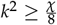. For a fixed value of *k*, the threshold speed given by Equation (C.5) is plotted in Figure S4. The above analysis and figure inform us of the following:

- For a finite selection strength *χ*_*α*_, *v*_*th*_ *< v*_*crit*_.
- As 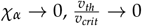.
- As 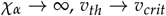.

Additionally, an expansion of Equation (C.5) for low selection strength and discarding higher-order terms reveals a power law dependence

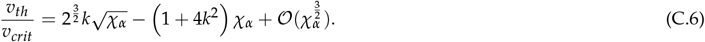

The polygenic simulations reveal a similar power-law relationship with an exponent that depends on wether the speed is scaled by the critical speed (*ωθ*) or by the square root of the equilibrium genetic variance (*σ*_*G*(*S*)_, see Figures 5c and S10). To explain how the two exponents are related, we recall that the equilibrium genetic variance can be approximated using the stochastic House-of-Card model (Bürger 2000) as

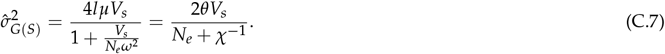

Now suppose that the threshold speed scaled by *ωθ* exhibits a power-law relationship with the selection strength *χ* as depicted in Figure 5c,

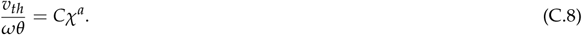

By scaling the threshold speed with 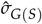, we get

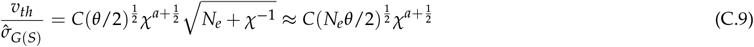

For 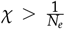, which is the relevant regime for our simulations. Thus the two exponents differ by ^1^, which is consistent with the numerical results.

## Supplementary Figures

**Figure S1.**
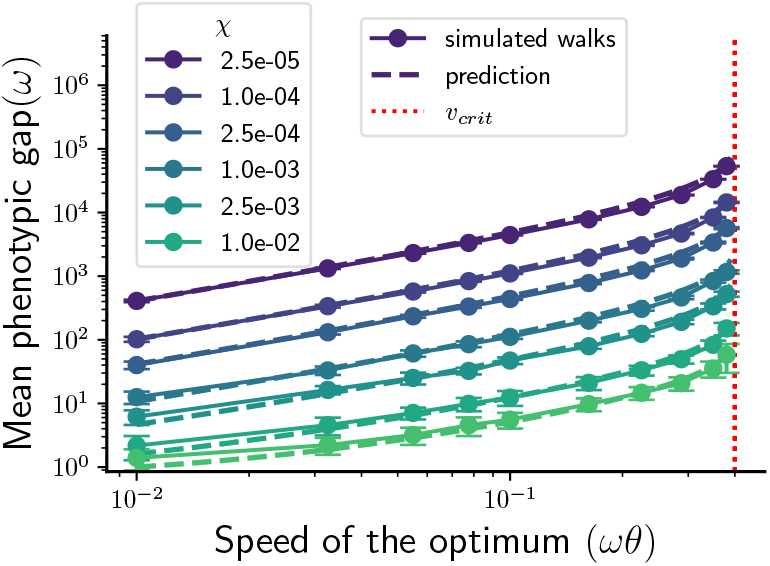
The mean phenotypic gap for adaptive walks as a function of the speed of the optimum.

**Figure S2.**
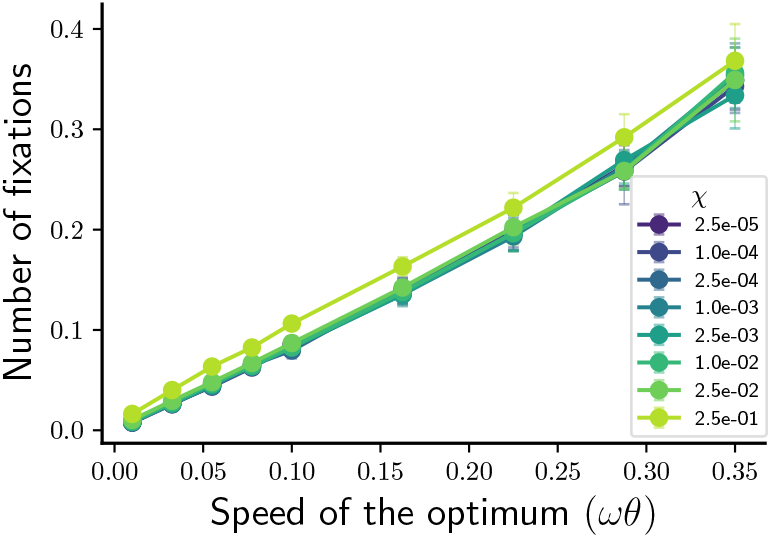
Number of fixations per generation in the stationary state as a function of the speed of the optimum for adaptive walks.

**Figure S3.**
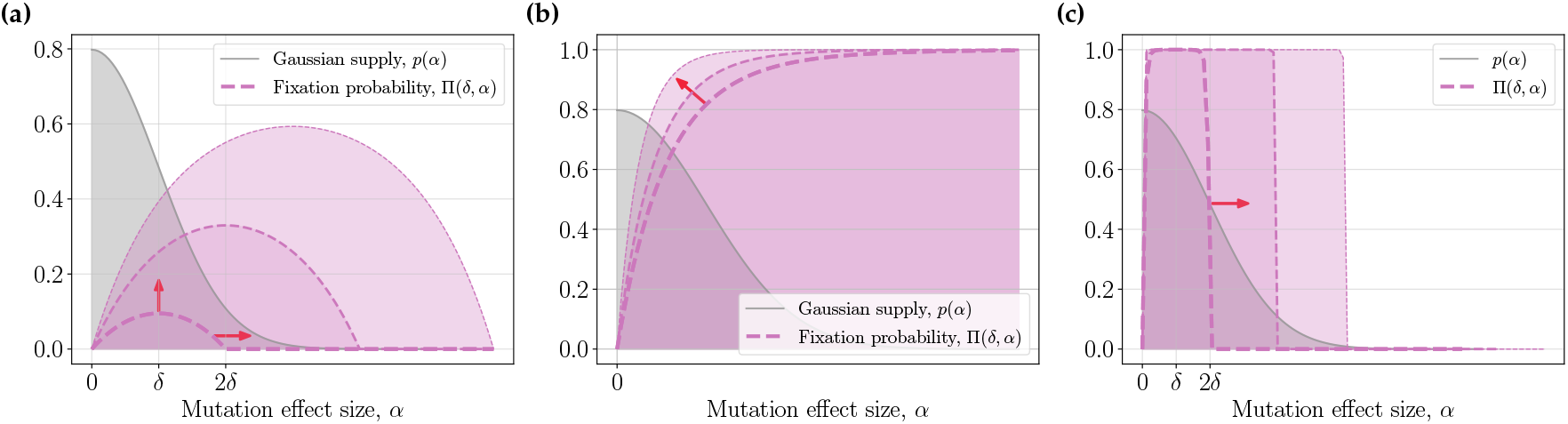
**(a)** The fixation probability and its support increase with increasing phenotypic gap (indicated by the red arrows). **(b)** At large gaps, while the support of the fixation probability keeps increasing, the fixation probability itself changes very little. The only observed change is the decreasing curvature at its maximum which increases the bias towards small effect mutations. **(c)** In the limiting case of high selection strength, the fixation probability is like a square function which spans increasingly more area of the supply with increase in the phenotypic gap.

**Figure S4.**
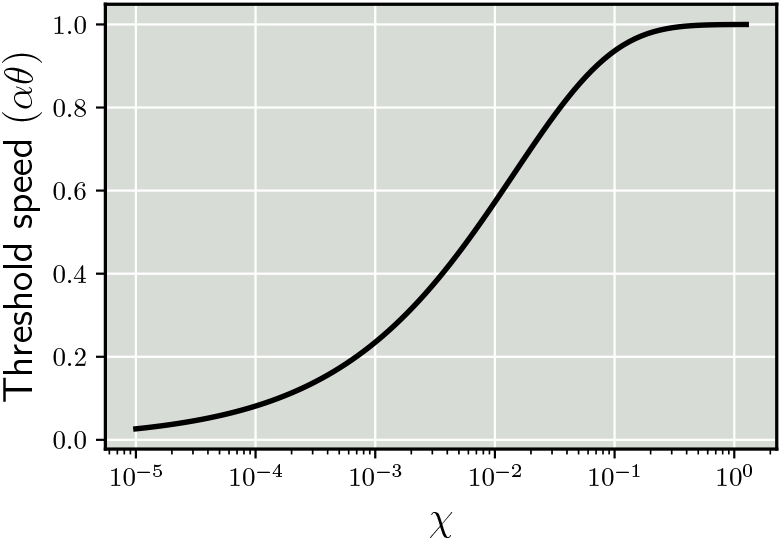
The threshold speed for a fixed value of the fitness threshold as predicted by Equation (C.5). The threshold speed increases with increasing trait parameter values and eventually converges to the critical speed.

**Figure S5.**
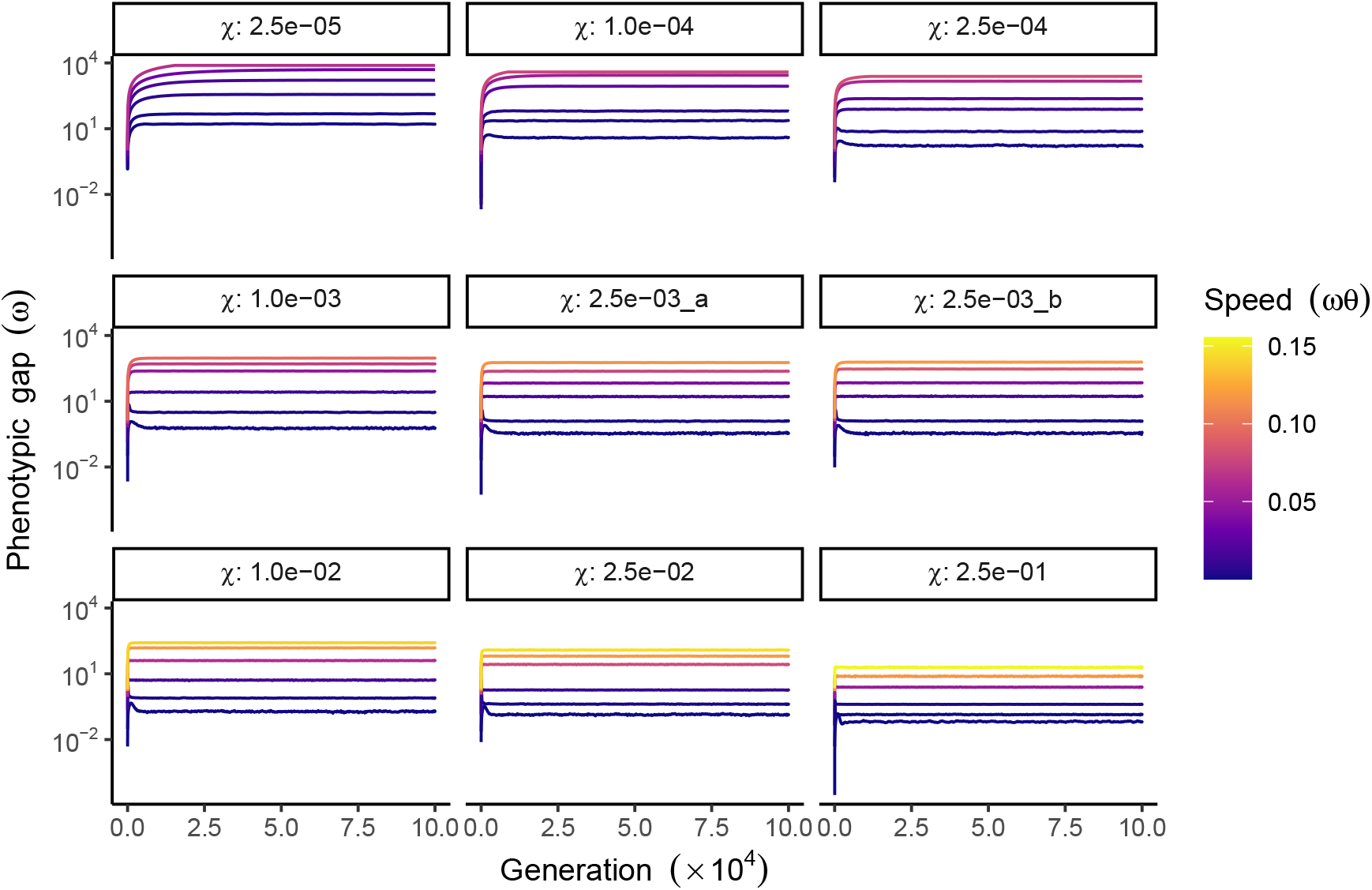
The phenotypic gap over time during optimum shifts in polygenic simulations. The mean phenotypic gap across 100 independent simulations during the optimum shift (generation 1-10 N). Each panel corresponds to a specific value of the selection strength *χ*, where ‘2.5e-03_a’ represents the parameter combination: *ω* = 0.05, *V*_*s*_ = 1, and ‘2.5e-03_b’ represents the combination:*ω* = 0.5, *V*_*s*_ = 100. The speed of the optimum shift is in units of *ωθ* and denoted by color gradients. The phenotypic gap is in units of *ω* and plotted on a logarithmic scale along the y-axis.

**Figure S6.**
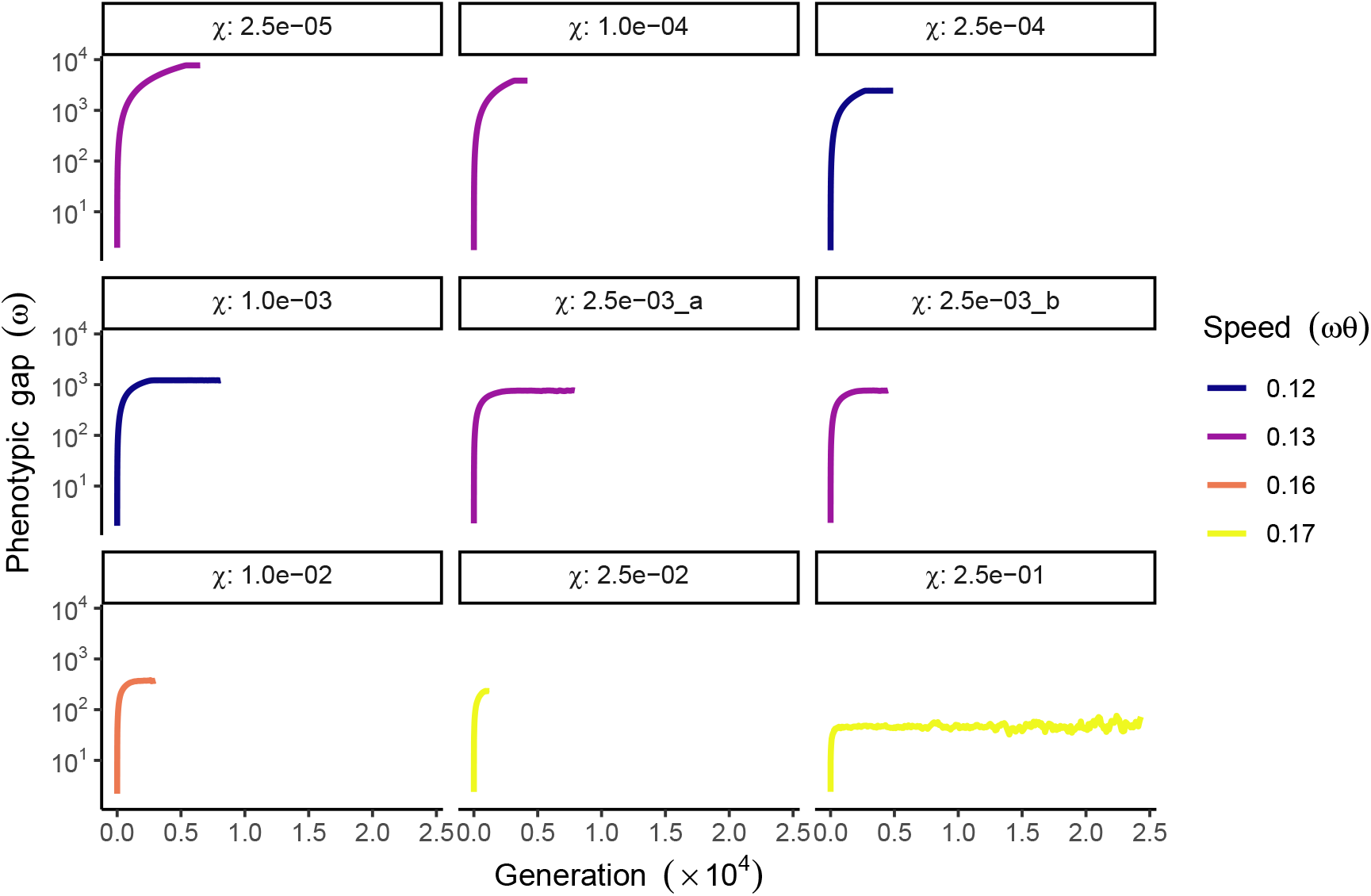
The phenotypic gap over time in extinct populations in polygenic simulations. The mean phenotypic gap across independent simulations (100 or less) from generation 1 to the simulation termination when all individuals’ fitness dropped to zero. Each panel corresponds to a specific value of *χ*, where ‘2.5e-03_a’ represents the parameter combination: *ω* = 0.05, *V*_*s*_ = 1, and ‘2.5e-03_b’ represents the combination: *ω* = 0.5, *V*_*s*_ = 100. The speed of the optimum shift is in units of *ωθ* and denoted by colors. The phenotypic gap is in units of *ω* and plotted on a logarithmic scale along the y-axis.

**Figure S7.**
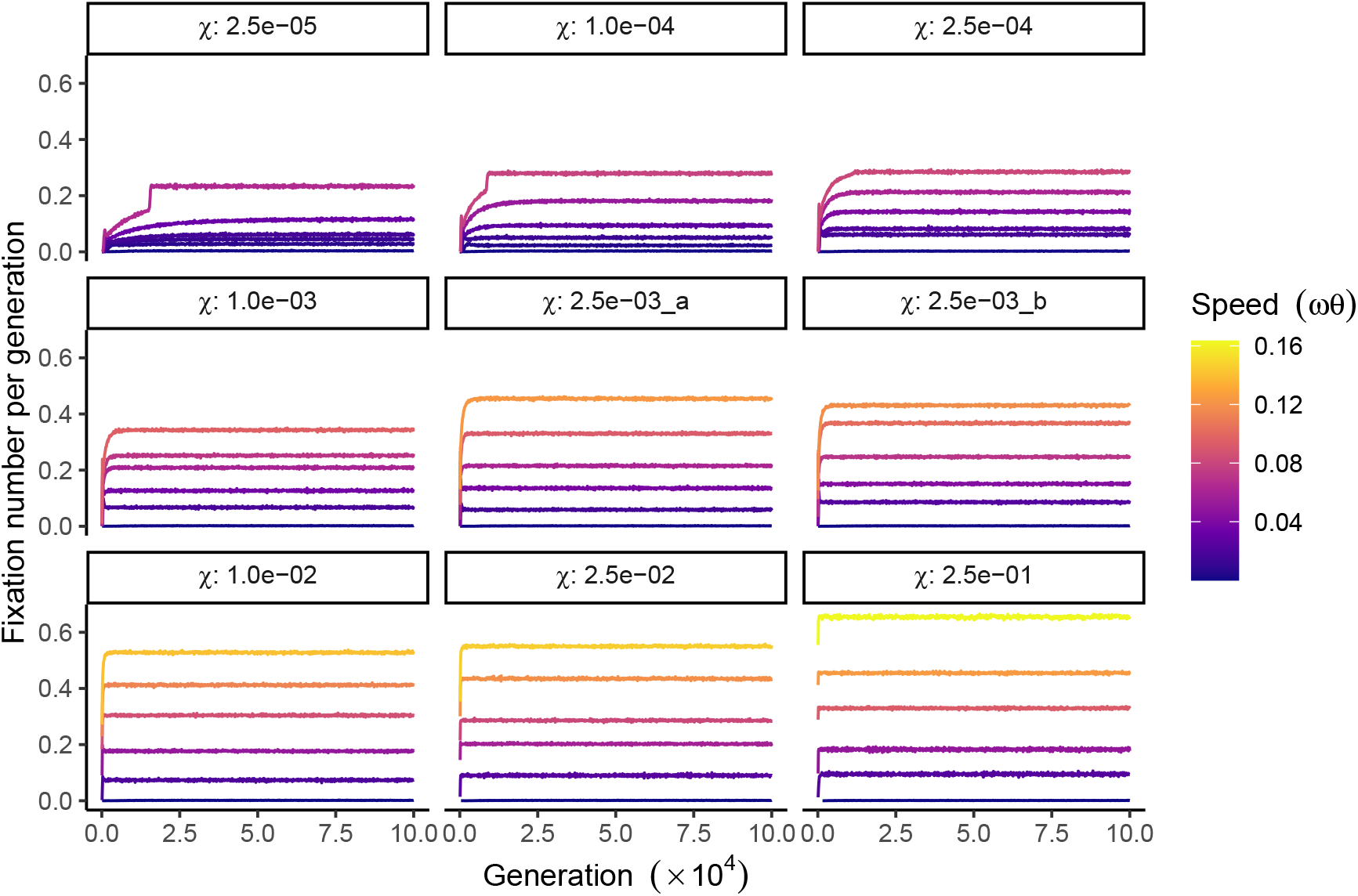
Mean number of mutation fixations in every generation in polygenic simulations. The y-axis represents the mean number of fixation events per generation (take an average every 100 generations) across 100 simulations during the optimum shift. Each panel represents a specific value of *χ*, where ‘2.5e-03_a’ represents the parameter combination: *ω* = 0.05, *V*_*s*_ = 1, and ‘2.5e-03_b’ represents the combination: *ω* = 0.5, *V*_*s*_ = 100. The speed of the optimum shift is in units of *ωθ* and denoted by color gradients.

**Figure S8.**
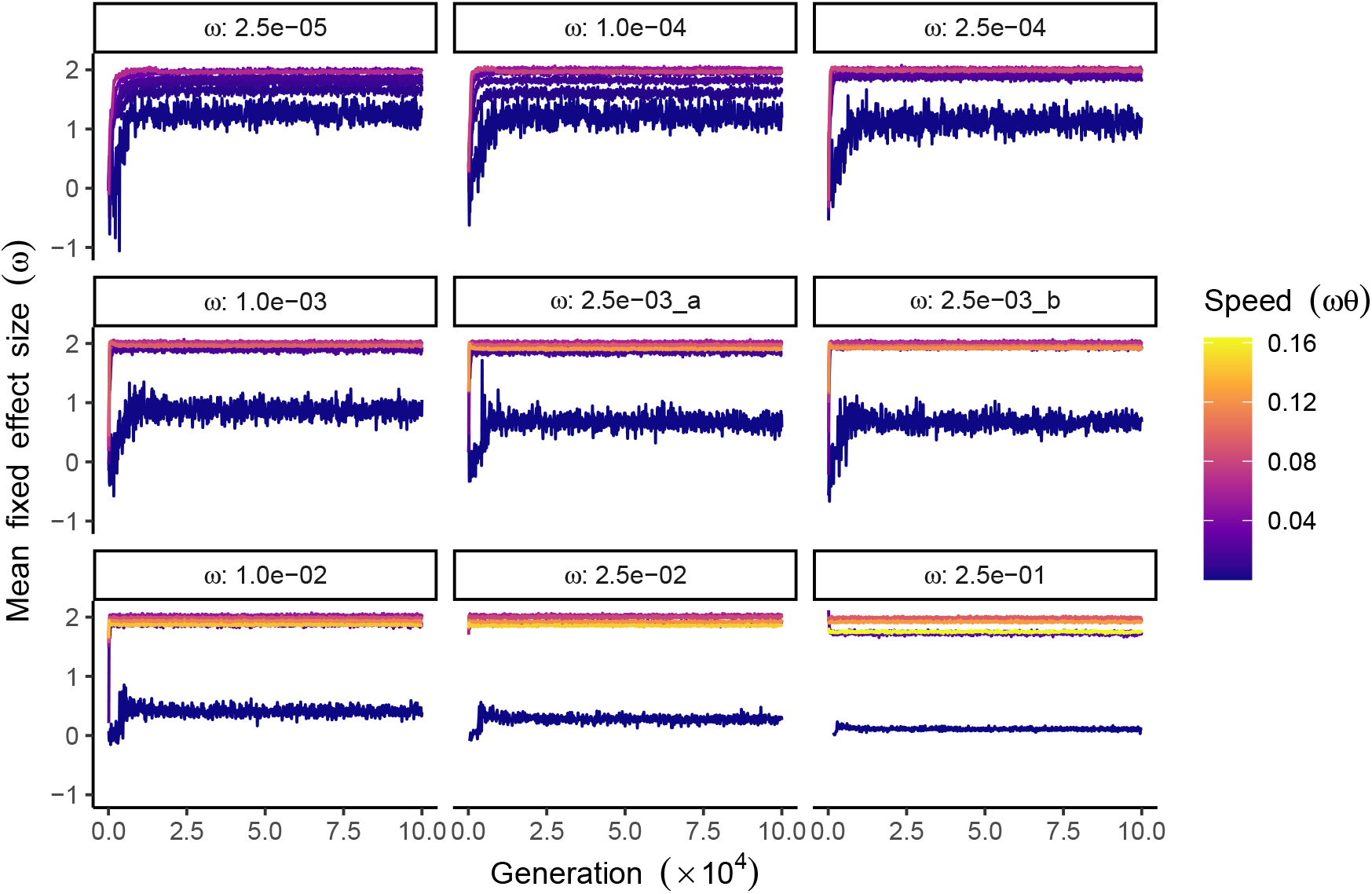
Mean effect size of fixed mutations every generation in polygenic simulations. The y-axis represents the mean effect size (in units of *ω*) of fixed mutations per generation (take an average every 100 generations) across 100 simulations during the optimum shift. Each panel represents a specific value of *χ*, where ‘2.5e-03_a’ represents the parameter combination: *ω* = 0.05, *V*_*s*_ = 1, and ‘2.5e-03_b’ represents the combination: *ω* = 0.5, *V*_*s*_ = 100. The speed of the optimum shift is in units of *ωθ* and denoted by color gradients.

**Figure S9.**
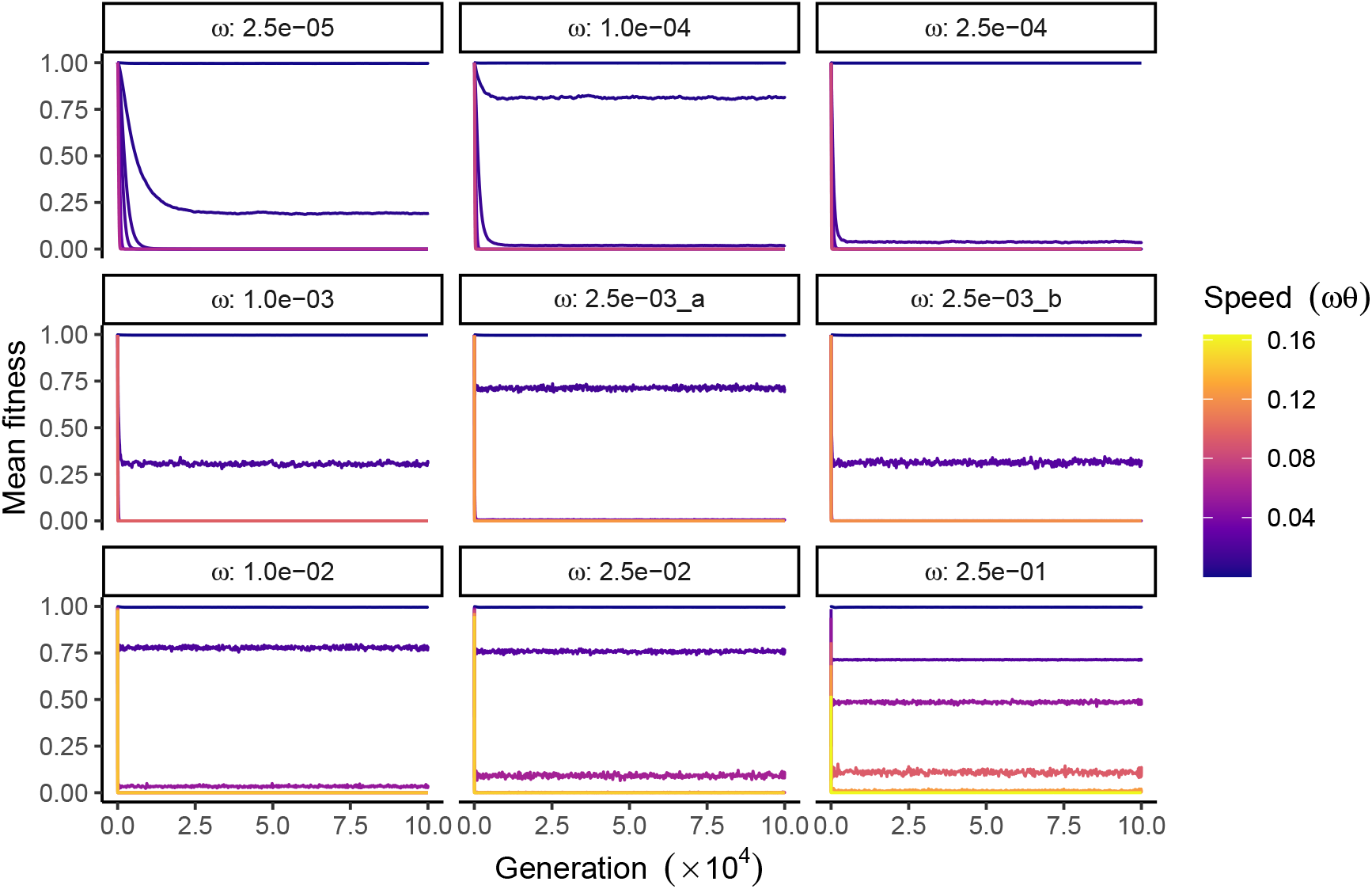
Mean fitness over time in polygenic simulations. The mean fitness of populations across 100 independent simulations during the optimum shift. Each panel corresponds to a specific value of *χ*, where ‘2.5e-03_a’ represents the parameter combination: *ω* = 0.05, *V*_*s*_ = 1, and ‘2.5e-03_b’ represents the combination: *ω* = 0.5, *V*_*s*_ = 100. The speed of the optimum shift is in units of *ωθ* and denoted by color gradients.

**Figure S10.**
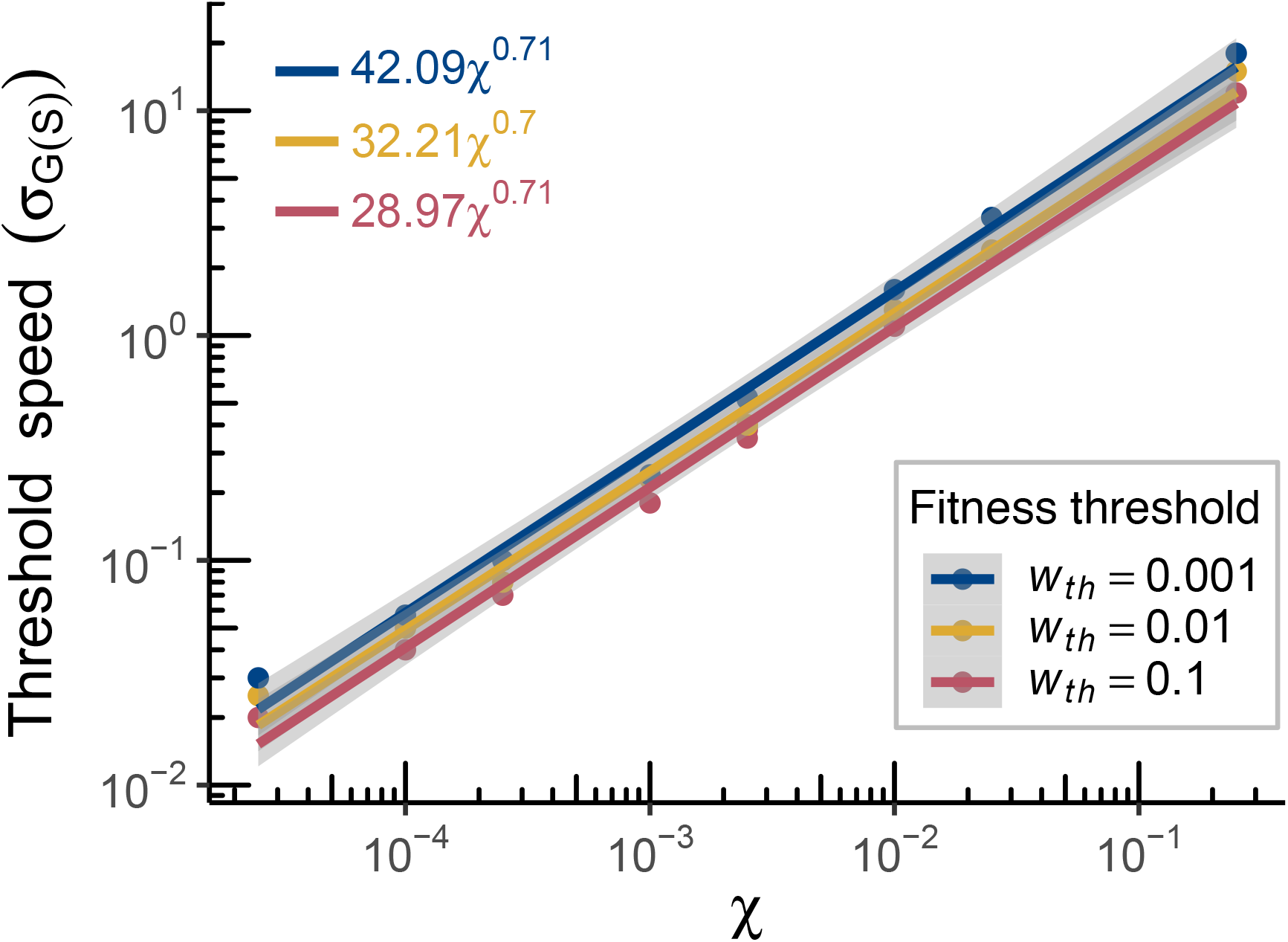
The threshold speed scaled by *σ*_*G*(*S*)_. In polygenic simulations, the threshold speeds, extracted at three fitness cutoffs (indicated by colors), are scaled by the square root of the equilibrium genetic variance under stabilizing selection (*σ*_*G*(*S*)_, calculated as the average genetic variance over the last 1000 generations before the optimum shift). The equations of the fitted lines are shown in the top-left corner. Both axes are shown on logarithmic scales. The relation to the power law relationship depicted in Figure 5c is explained in Equation (C.9).

**Figure S11.**
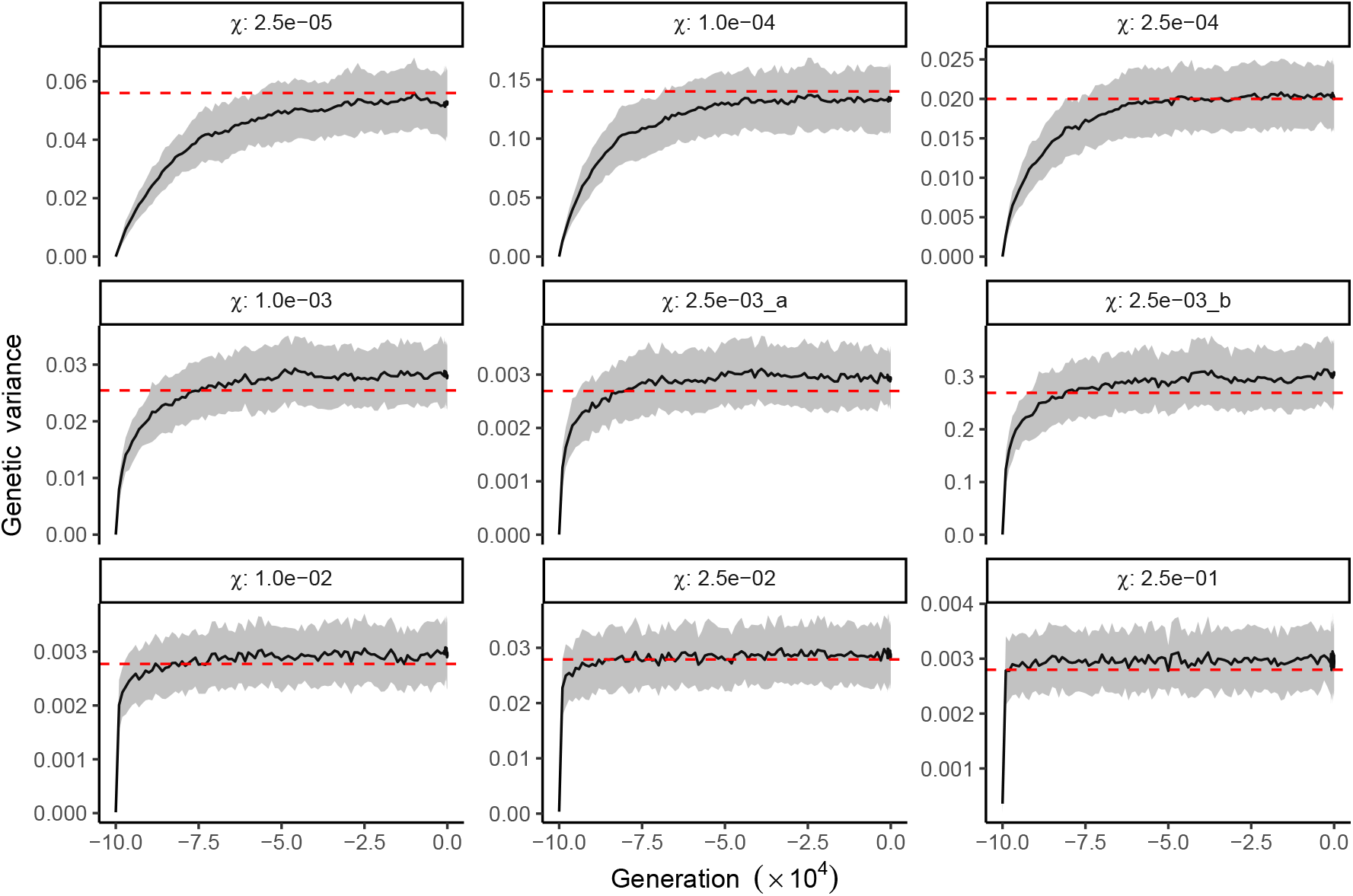
Mean genetic variance before optimum shifts in polygenic simulations. The black line represents the mean genetic variance of phenotype across 100 independent simulations before the optimum shift (generation -10 N - 0), with the surrounding grey area indicating the standard deviation of genetic variance across 100 replicates. The red dashed line denotes the theoretical genetic variance predicted by the stochastic House-of-Cards model in Equation (C.7) (Bürger 2000).

**Figure S12.**
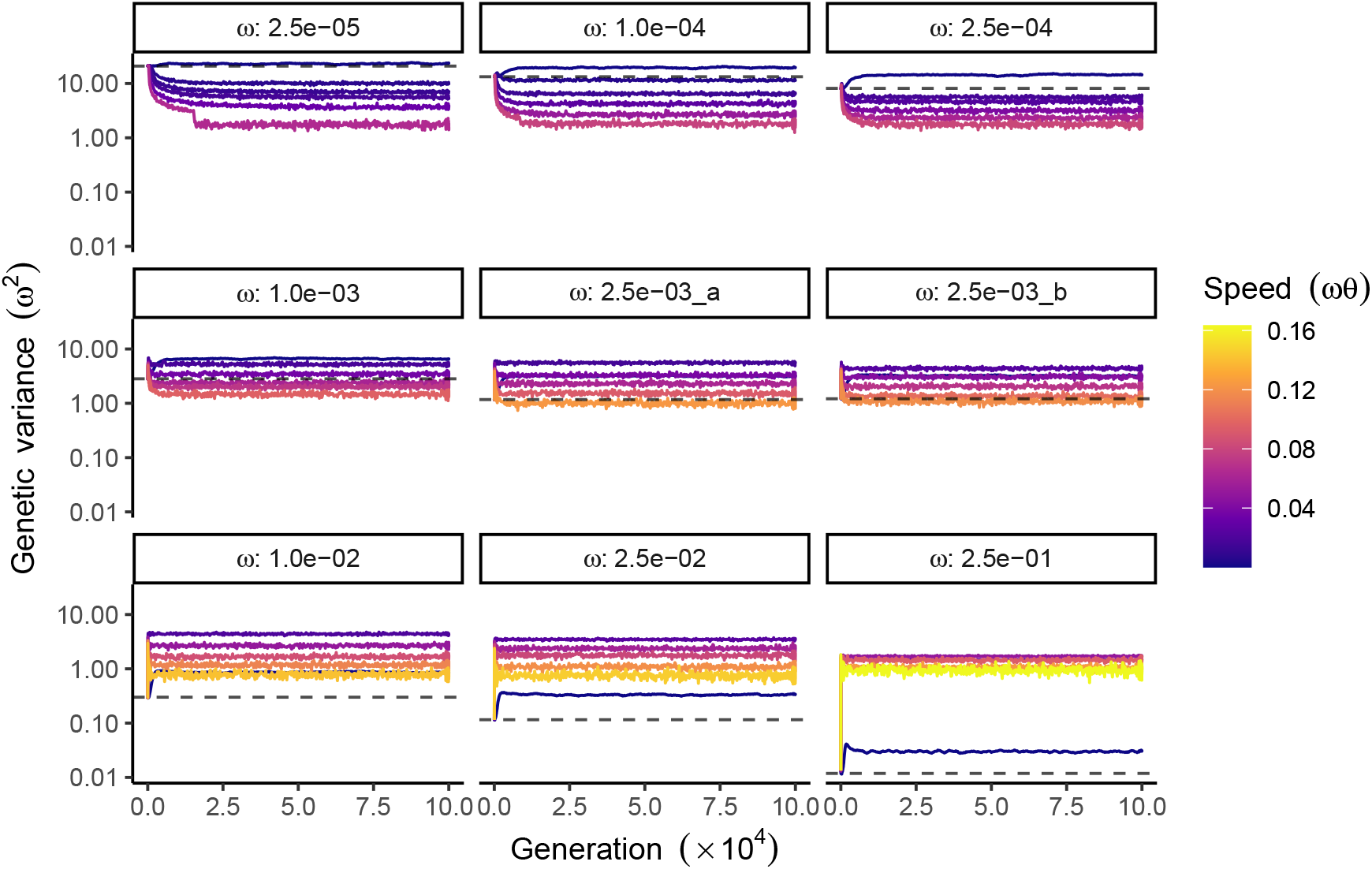
Mean genetic variance during optimum shift in polygenic simulations. The mean genetic variance of phenotype (in units of *ω*^2^) across 100 independent simulations during optimum shift (generations 1-10 N). Each panel corresponds to a specific value of *χ*, where ‘2.5e-03_a’ represents the parameter combination: *ω* = 0.05, *V*_*s*_ = 1, and ‘2.5e-03_b’ represents the combination: *ω* = 0.5, *V*_*s*_ = 100. The speed of the optimum shift is in units of *ωθ* and denoted by color gradients. The black dashed line denotes the average genetic variance over the last 1000 generations before the optimum shift 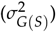.

**Figure S13.**
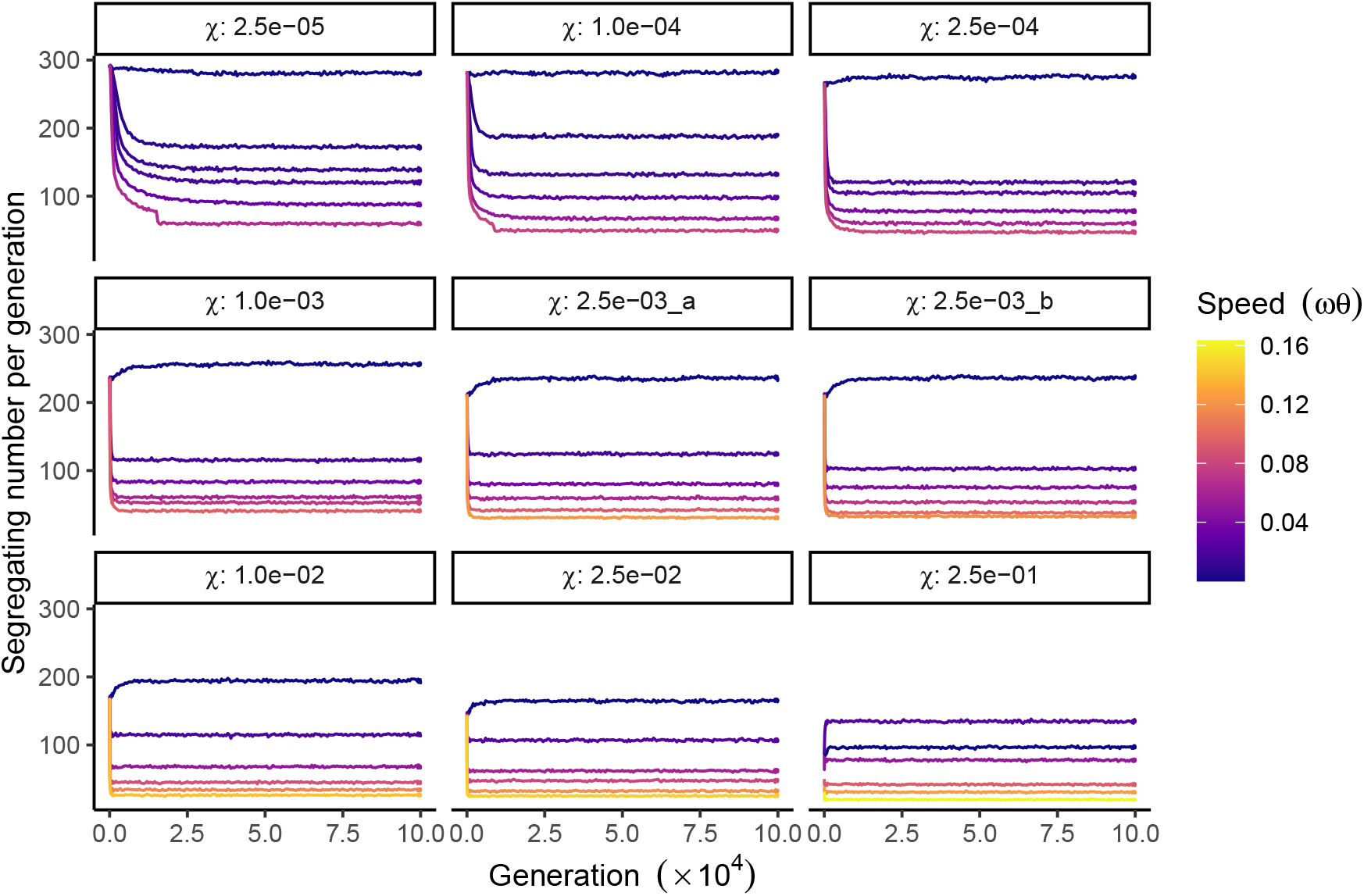
Mean number of segregating sites in polygenic simulations. The mean number of segregating sites per generation across 100 independent simulations during the optimum shift. Each panel corresponds to a specific value of *χ*, where ‘2.5e-03_a’ represents the parameter combination: *ω* = 0.05, *V*_*s*_ = 1, and ‘2.5e-03_b’ represents the combination: *ω* = 0.5, *V*_*s*_ = 100. The speed of the optimum shift is in units of *ωθ* and denoted by color gradients.

**Figure S14.**
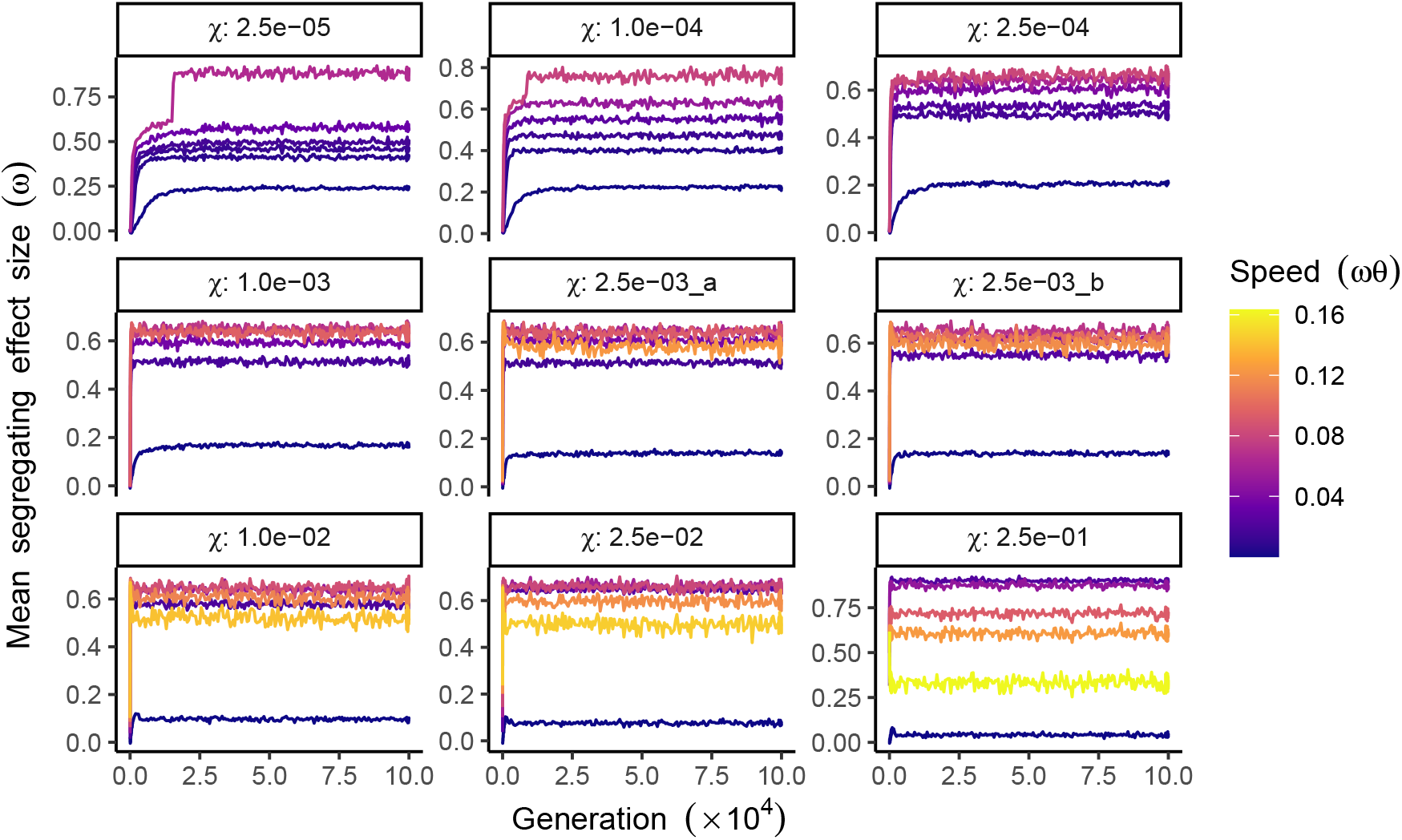
Mean effect size of segregating sites in polygenic simulations. The mean effect size of segregating sites (in units of *ω*) per generation across 100 independent simulations during the optimum shift. Each panel corresponds to a specific value of *χ*, where ‘2.5e-03_a’ represents the parameter combination: *ω* = 0.05, *V*_*s*_ = 1, and ‘2.5e-03_b’ represents the combination: *ω* = 0.5, *V*_*s*_ = 100. The speed of the optimum shift is in units of *ωθ* and denoted by color gradients.

**Figure S15.**
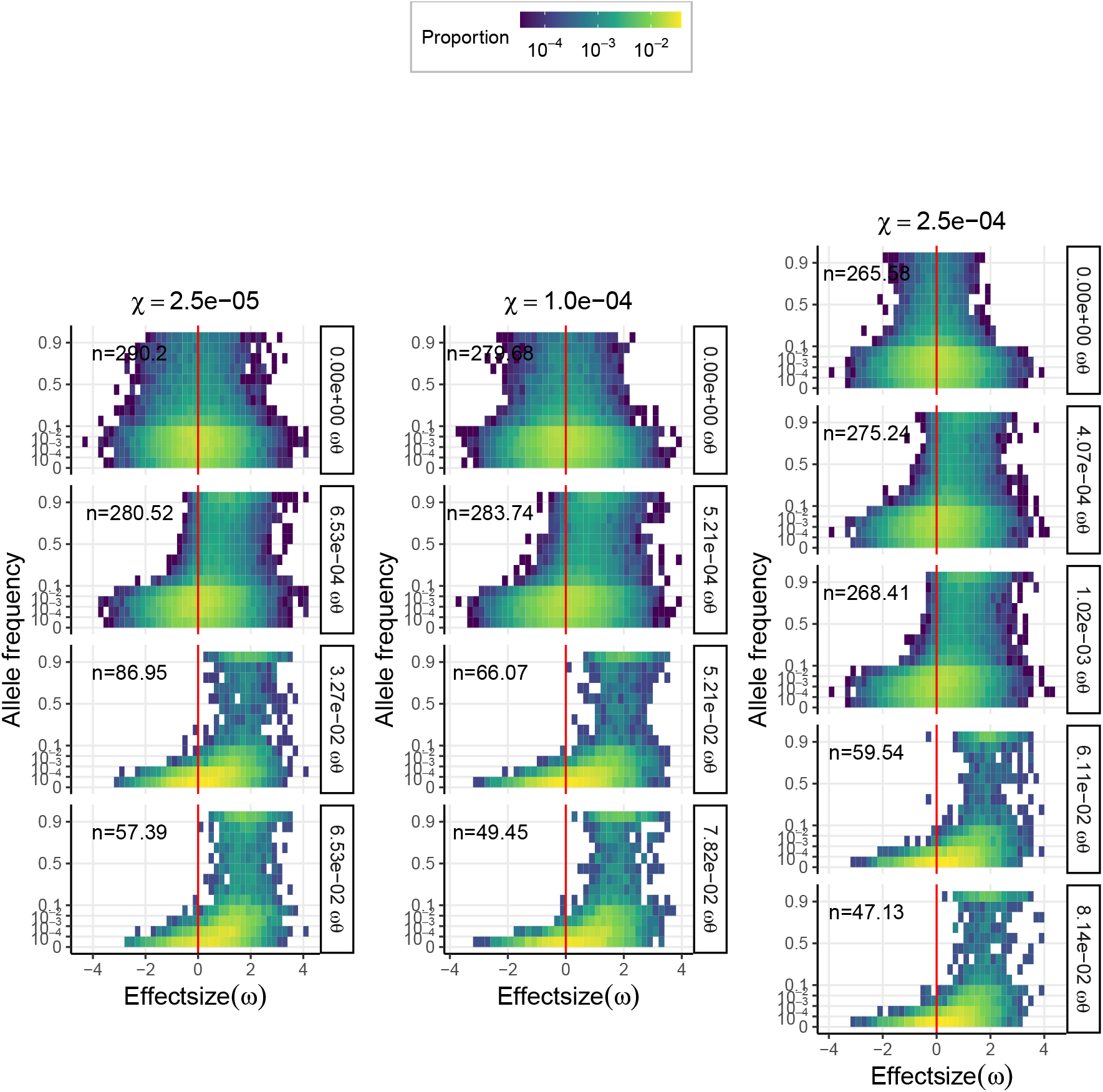
Genetic architecture in different scenarios. Genetic architecture matrices at generation 10 N for *χ* = 2.5 × 10^−5^, 1.0 × 10^−4^ and 2.5 × 10^−4^. Effect size bins range from negative to positive values and are centered at zero (red line). The facet labels on the right of the panels indicate the speeds of optimum shift (in units of *ωθ*). The color represents the proportion of segregating mutations within a given range of effect sizes and allelic frequencies. The number shown in each panel denotes the average number of segregating sites across 100 independent simulations.

**Figure S16.**
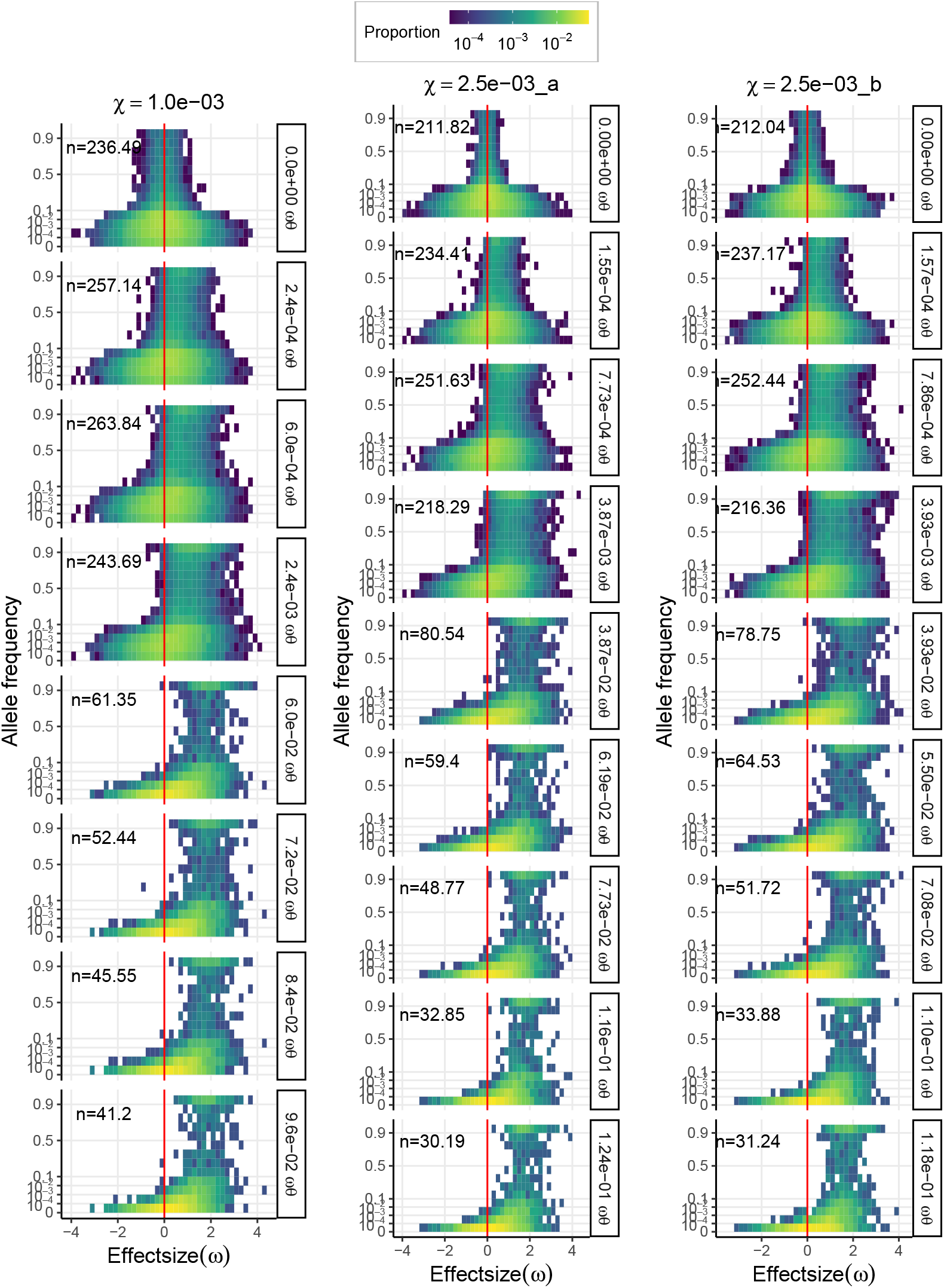
Genetic architecture in different scenarios. Genetic architecture matrices at generation 10 N for *χ* = 1 × 10^−3^ and 2.5 × 10^−3^, where ‘2.5e-03_a’ represents *ω* = 0.05, *V*_*s*_ = 1 and ‘2.5e-03_b’ represents *ω* = 0.5, *V*_*s*_ = 100. Other figure parameters are the same as in Figure S15.

**Figure S17.**
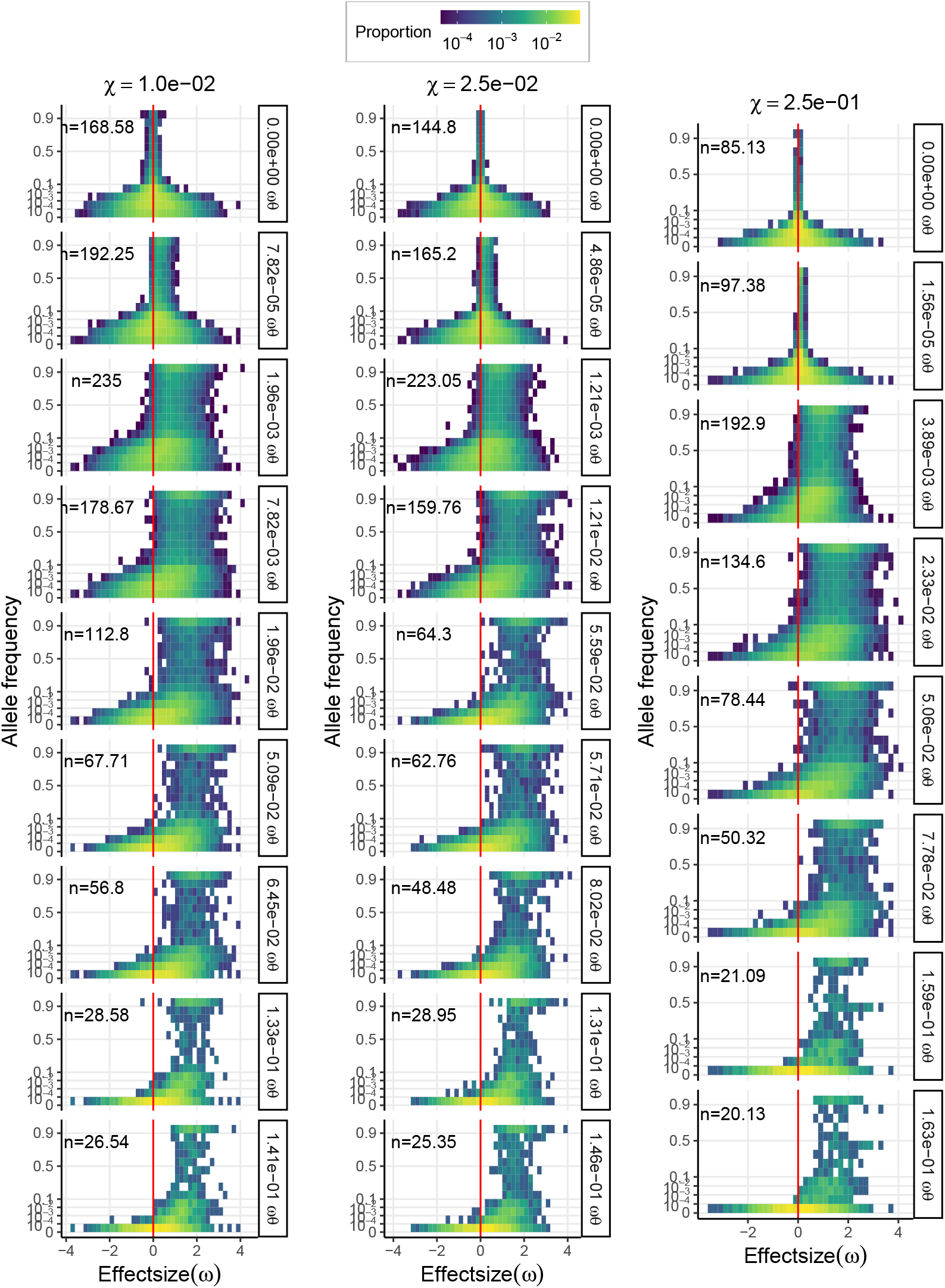
Genetic architecture in different scenarios. Genetic architecture matrices at generation 10 N for *χ* = 1 × 10^−2^, 2.5 × 10^−2^ and 2.5 × 10^−1^. Other figure parameters are the same as in Figure S15.

**Figure S18.**
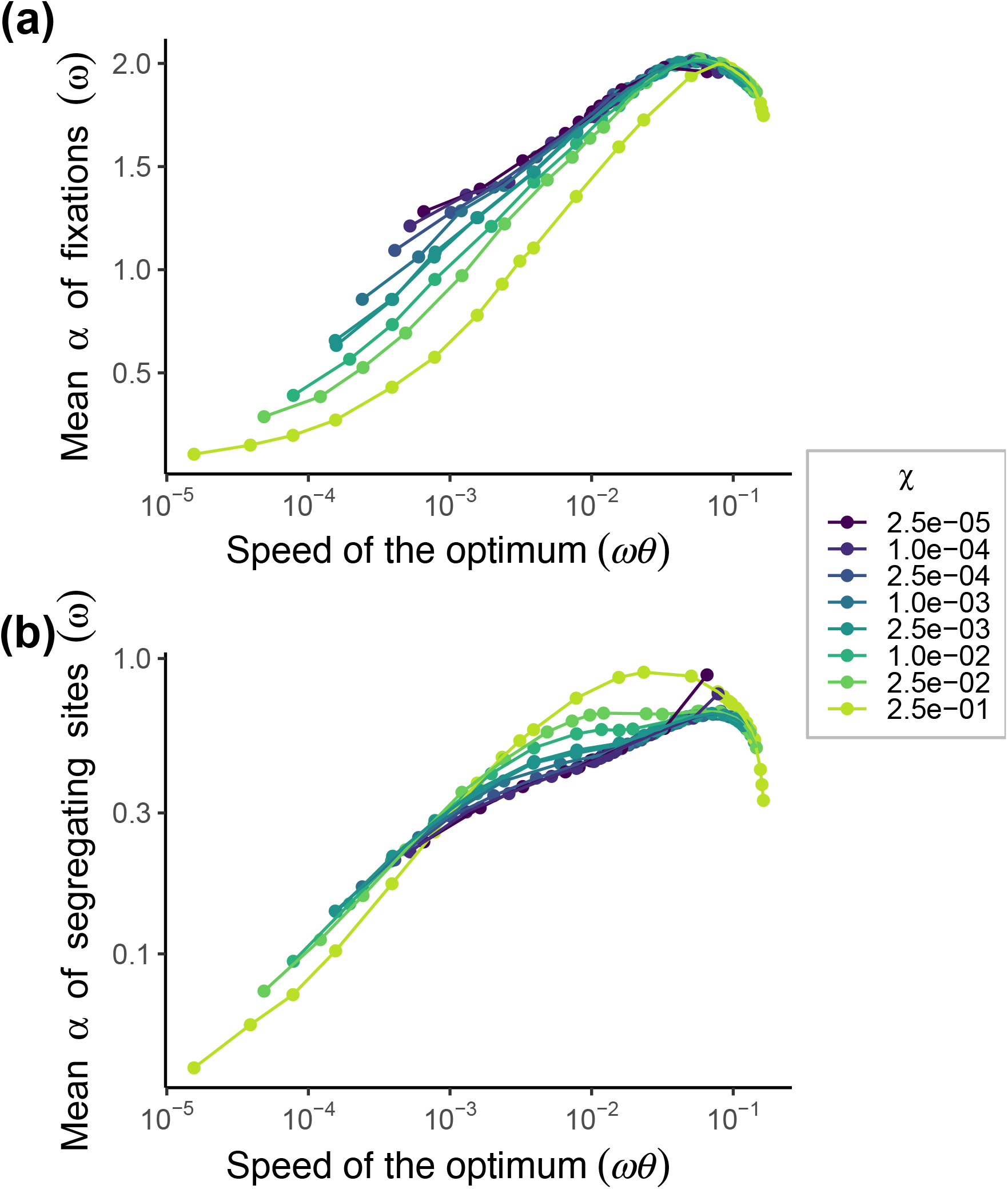
The comparison between the mean effect size of fixations and segregating sites in polygenic simulations. The mean effect sizes of fixed mutations (a) and segregating sites (b) are in units of *ω*, averaged over the last 1000 generations of 100 independent simulations. The x-axis is on a log scale.

**Figure S19.**
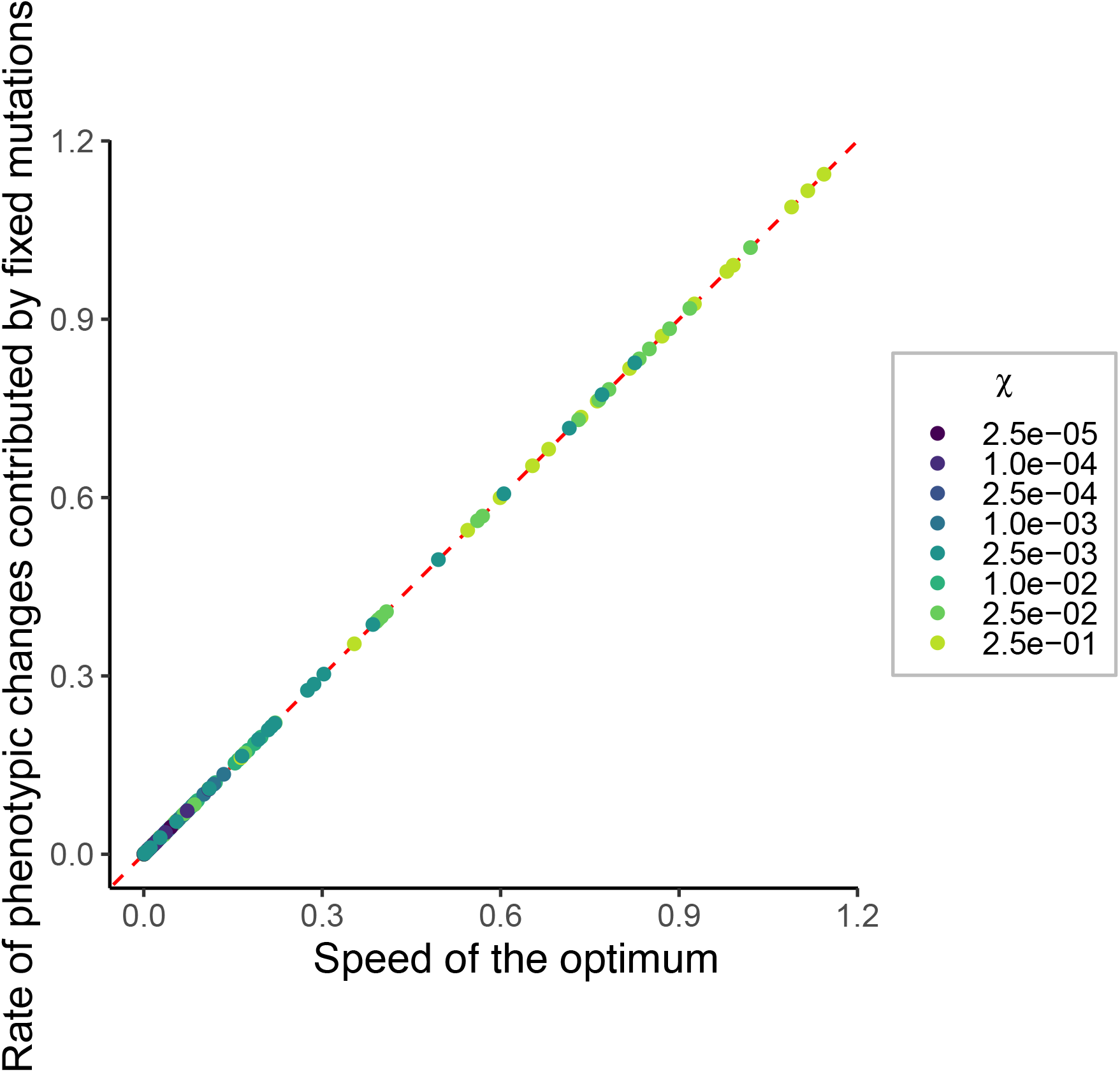
The rate of phenotypic changes contributed by mutation fixations is the same as the speed of optimum shift. The rate of phenotypic changes contributed by mutation fixations is calculated by the formula 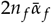, where *n_f_* (number of fixation events per generation) and 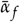 (mean effect size of fixed mutations) correspond to stationary-state measurements in Figures 3a and 3b, respectively. The red diagonal (*y* = *x*) denotes perfect agreement between the rate of phenotypic changes contributed by fixed mutations and shift speed.

**Figure S20.**
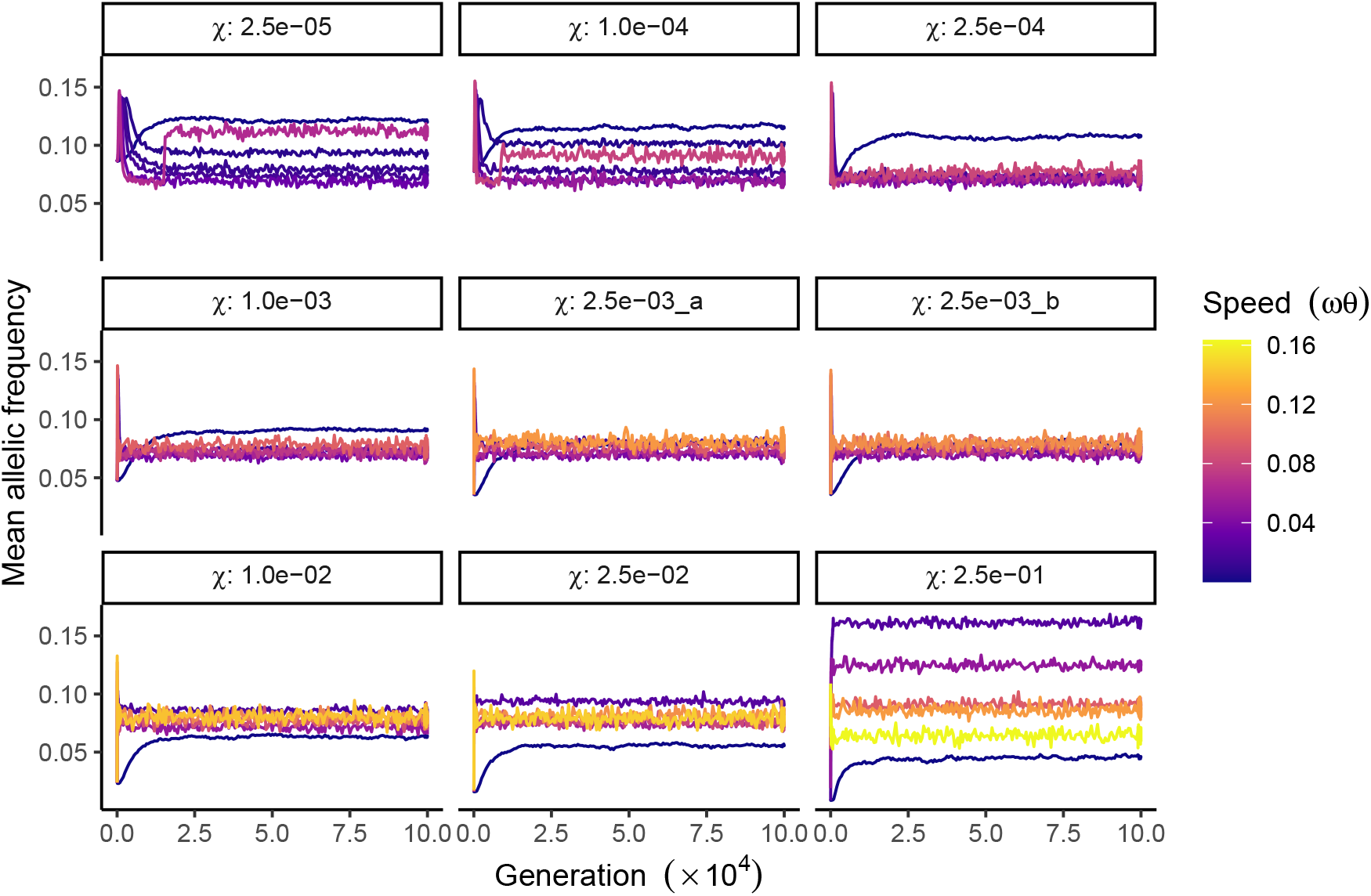
Mean allelic frequency in polygenic simulations. The mean allelic frequency of segregating sites across 100 independent simulations during the optimum shift. Each panel corresponds to a specific value of *χ*, where ‘2.5e-03_a’ represents the parameter combination: *ω* = 0.05, *V*_*s*_ = 1, and ‘2.5e-03_b’ represents the combination: *ω* = 0.5, *V*_*s*_ = 100. The speed of the optimum shift is in units of *ωθ* and denoted by color gradients.

## Footnotes

1 The dependence on time will be suppressed from hereon since we will only consider instances when a mutation occurs.

2 ⟨.⟩_*X*_ indicates average with respect to the random variable *X*

## Notes

### Competing Interest Statement

The authors have declared no competing interest.

